# Cholesterol and p53 promote senescence and systemic fibrosis in metabolic dysfunction-associated steatohepatitis

**DOI:** 10.64898/2026.09.02.748892

**Authors:** Celine I Wittke, Dale M Watt, Liam Butler, Anabel Martinez Lyons, Cassie J Clarke, Clara Mullen, Nicola Clements, Amy Lawlor, Dimitris Athineos, Ashleigh Young, Douglas Strathdee, Colin Nixon, Leonard J Nelson, Thomas G Bird, Karen H Vousden, Karen Blyth, Jennifer Crowe, Timothy J Humpton

## Abstract

**Background & aims:** TP53 (p53) coordinates diverse cellular stress response programmes including pro-survival activities, senescence, and cell death. During tissue damage, p53 can shape both the local cellular response to injury, including the fibrotic response, and influence distal organ biology. Fibrosis in the liver is a major driver of hepatocellular carcinoma (HCC) risk within metabolic dysfunction-associated steatohepatitis (MASH). It is also an important determinant of dysfunction in multiple distal tissues including the kidneys, lungs, and heart. Despite significant clinical burden, our understanding of the molecular determinants of fibrotic MASH and its relationship to multiorgan fibrosis remain incomplete. Here, we investigate local and systemic effects of hepatocellular p53 activity and cholesterol during MASH development, with implications for disease prevention.

**Methods:** This study utilised a genetic model of stabilised p53, diet-induced MASH models with varying cholesterol compositions, and an *in vitro* obesogenic system to investigate p53 activity during liver disease development. Non-invasive imaging and histopathological analyses were employed to monitor p53 activity, MASH, and multiorgan fibrosis *in vivo*. Complementary approaches, including *in vitro* human multicomponent liver spheroids, cytokine arrays, and analyses of human MASH transcriptomic and proteomic datasets, were used to examine molecular drivers and patient relevance.

**Results:** Using an inducible mouse model of MDM2 E3 ubiquitin ligase deficiency to stabilise p53, we report that hepatocellular MDM2 E3 loss results in progressive fibrotic damage, robust hepatocellular expression of the p53 target gene *CDKN1A/p21* (p21),and induces p21 and fibrosis in the kidneys of male mice in a sex-specific manner. In diet-induced MASH, we observe cholesterol and p53-dependent development of liver fibrosis, high expression of hepatocellular p21, and induction of p21 and fibrosis in the kidneys of male mice—reminiscent of features observed in MDM2 E3-deficient mice. We also observe fibrosis in the lungs and heart of male MASH mice. Both a cholesterol-free obesogenic diet and liver-specific loss of p53 mitigate hepatic fibrosis and systemic induction of p21 and fibrosis. Mechanistically, p53 induces hepatic expression of senescence-associated secretory phenotype (SASP) factors, including GDF15, *in vivo*. A human multicomponent LiverACE spheroid model showed a concordant trend towards increased GDF15 protein abundance under steatotic stress, while in humans, elevated circulating GDF15 levels in advanced MASH correlate with increased TNFRSF1A and EPHA2, circulating markers linked to kidney injury.

**Conclusions:** Our work identifies undue p53 activity within the liver as a driver of multiorgan fibrosis in a sex-specific manner, affecting male but not female mice. We implicate cholesterol in promoting this pro-fibrotic environment *in vivo* and highlight circulating factors that could identify at-risk patients for multiorgan fibrosis in MASH.

**IMPACT AND IMPLICATIONS:** P53 is a potent tumour suppressor that is frequently mutated or lost in HCC. Our findings suggest that dietary cholesterol promotes detrimental overactivation of p53 in MASH and facilitates fibrosis within the liver and distal organs. Further studies evaluating the efficacy of targeting hepatic p53 in combination with dietary or weight loss interventions could provide new treatment approaches for systemic fibrosis and MASH.

## INTRODUCTION

Metabolic dysfunction-associated steatohepatitis (MASH) is a prevalent and insidious liver condition with few treatment options and is a known precursor to primary liver cancer^1^. MASH-derived liver cancer (Hepatocellular Carcinoma / HCC) is recalcitrant to existing standard-of-care therapies as well as novel immunotherapies^2,3^. Liver fibrosis in MASH is not only a strong risk factor for MASH-HCC^4–6^ but has also been linked to increased risk of chronic kidney disease (CKD)^7^, respiratory disease^8,9^, and adverse outcomes in cardiovascular disease (CVD)^10–12^. Within this context, the development of anti-fibrotic interventions against MASH, exemplified by the success of the thyroid hormone β receptor agonist Resmetirom and the GLP-1 agonist Semaglutide for liver scarring, are encouraging^13,14^. However, due to the high cost, limited availability, and severe side-effects associated with these treatments, there remains a pressing need for greater understanding of the molecular drivers of liver fibrosis to spur development of novel anti-fibrotic approaches^15^.

Diet composition has important implications for the onset and progression of metabolic dysfunction-associated steatotic liver disease (MASLD) to MASH due to local and systemic effects^16^. Dietary cholesterol, for example, has been implicated as a substantial accelerant for MASH development in murine models and humans^17–22^. MASLD to MASH progression is also strongly linked with features of metabolic syndrome^23–26^. These, in turn, promote fibrotic MASH, especially in the context of type 2 diabetes mellitus (T2DM)^27^. Metabolic syndrome and T2DM similarly exacerbate fibrosis in organs distal to the liver including within the kidneys (CKD), the lungs (as in idiopathic pulmonary fibrosis (IPF)), and the heart (CVD)^28–30^. Of particular concern, patients with multiorgan fibrosis burden harbour significantly increased mortality risk compared to those with low fibrosis burden^31^. Multiorgan morbidity is not limited to MASH amongst liver pathologies. Severe acute liver damage has also been shown to trigger dysfunction of distal organs *in vivo*—with implications for patients diagnosed with acute indeterminate hepatitis^32^. In this context, loss of control over the TP53 (p53) pathway promotes hepatocellular senescence, a state of permanent cell cycle arrest^33^, that in turn causes multiorgan senescence that is directed by the release of senescence-associated secretory phenotype (SASP) factors^32^. These findings, alongside positive correlations between fibrotic MASH and CKD, CVD, and IPF, suggest a liver-centric driver for multiorgan fibrotic pathology^7–12^.

The tumour suppressor p53 is reported to exert disparate effects on MASH disease trajectory. On the one hand, increased expression of either *TP53* or the p53 target gene *CDKN1A/p21* (p21) is correlated with MASH, liver fibrosis, and T2DM^34,35^. High hepatocellular p21 expression in patients with acute indeterminate hepatitis is also predictive of multiorgan failure^32^. On the other hand, moderate activation of p53 is known to limit inflammation, reduce fibrosis, and protect against fatty-acid induced apoptosis in the context of acute liver injury and provides anti-MASH protection through enhanced redox control^36–42^. These disparate outcomes likely reflect context-specific balancing of p53 activity between pro-survival pathways, senescence, and cell death^43–45^. For example, protective p53 activation can be achieved with little observed hepatic p53 stabilisation and reduced p53 binding affinity for some target genes^46–48^. Abundant stabilisation of hepatic p53, in contrast, is a potent driver of senescence and cell death that can further promote multiorgan dysfunction^32^.

Our work identifies chronic elevated p53 activity within the liver as a mediator of fibrotic liver damage, advanced MASH, and concomitant multiorgan fibrosis. We also implicate cholesterol and SASP factors including GDF15 in promoting this environment. Using extant human data from MASH patients and *in vitro* hepatocellular models, we further highlight a relationship between circulating GDF15 and markers of kidney damage in MASH patients that could identify an at-risk group for multiorgan fibrosis. Further studies evaluating the efficacy of targeting hepatic p53 alongside diet or weight loss interventions to prevent systemic fibrosis could provide new treatment approaches for multiorgan fibrosis and MASH.

## MATERIALS AND METHODS

### Mice

Procedures involving mice were performed under Home Office licence numbers PP6345023 and PP1179816. The programme of work was sanctioned by Local Ethical Review Process (University of Glasgow) and conducted in accordance with the Animals (Scientific Procedures) Act 1986. Mice were housed on a 12-hour light/12-hour dark cycle. They were provided with diet and water *ad libitum* and given environmental enrichment in the form of Sizzle-Nest bedding or Happi Mats Nesting Squares (IPS) and polycarbonate tunnels. To reduce animal anxiety and improve welfare, non-aversive handling using polycarbonate tunnels was utilised throughout each experiment.

*p53^FL/FL^* (*Trp53*^tm1Brn^), PG13-iRFP p53 reporter (*Hprt^Tm1(p53RE-iRFP^ ^[pg13])Bea^*), and *Mdm2^Ex5/6Δ^*(*Mdm2*^tm2.1Glo^) mice have been described previously^48–50^ and were backcrossed to C57BL/6J (N10). Cohort p53 mice used in Figures 3-5 were created by interbreeding *p53^FL/FL^* and PG13-iRFP mice. The creation of both Tm1 and Tm1.1 *Mdm2^I438K^* mice has been previously described and included interbreeding with *Rosa*- Cre^ER^ [Gt(ROSA)26Sor^tm2(cre/ERT2)Brn^] mice as previously described for the purposes of whole-body *Mdm2^I438K^* induction in Figure S1^51^, although functional validation of Tm1.1 *Mdm2^I438K^* mice has not been reported. For further information on whole-body *Mdm2^I438K^* induction, see supplementary methods.

Within experimental cohorts, mice were age and littermate matched as much as possible. Treatment with AAV8-TBG-Cre (AAV8.TBG.PI.Cre.rBG, Addgene, 107787-AAV8) or AAV8-TBG-null (AAV8.TBG.PI.Null.bGH, Addgene, 105536-AAV8) was performed as previously described^52^. Based on initial diet results that indicated more pronounced hepatocellular p53 activity and more advanced liver disease in males on the high fat high sugar high cholesterol diet than females, supporting studies using the high fat high sugar + cholesterol diet (Figure S4) were examined in only male cohorts. All other mouse experiments included analysis of both male and female mice. For the purposes of this study, clinical endpoint was defined as a 20% weight loss from baseline according to the relevant protocols of the Home Office Licenses governing experimental work.

Due to differences in weight gain amongst experimental mice and the differing physical appearance of high fat diet formulations compared to chow, it was not possible to blind researchers to diet treatments. However, researchers were blinded to the genotype of each animal throughout longitudinal analyses and during sample collection. In the downstream analyses of experiments involving mice, quantifications were conducted on a random order of samples. Sample cohort information was blinded until the final summation of results.

### Diet experiments

60-75-day old male and female chow-fed mice were shifted onto either a high-fat and high-sugar cholesterol-free (HFHS) diet, a high fat high sugar high cholesterol (HFHSHC) diet, the HFHS diet reformulated to contain cholesterol (HFHS+Chol), or left on mouse chow (control diet, DS801752G10R, Special Diet Services). HFHS diet (TestDiet 58R3), HFHSHC diet (TestDiet 5ZSF), and HFHS+Chol diet (TestDiet 5ZY4) were purchased from TestDiet. For further information on diet composition, see supplementary methods.

### In vivo imaging

Longitudinal imaging was conducted as previously described^42^ using a Pearl Impulse Small Animal Imaging System (LI-COR). All mice were scanned at a resolution of 85 µm in the 700 nm channel only. Scan images in the figures are presented with false colour LUTs which were held constant across the images shown. For quantification, Image Studio software (LI-COR, V5.5) was used to identify the p53 reporter (p53rep) signal within the liver-region of each mouse. These measurements were normalised to the baseline liver- region signal for each mouse, recorded 7 days after diet change.

### *Ex vivo* imaging of tissues

Harvested tissues were fixed, orientated, and imaged as previously described using the same scan settings as for *in vivo* imaging^42^. Scans were analysed and presented as for *in vivo* imaging.

For further information on experiments involving mice, including strain background analyses, see supplementary methods.

### Histology, Immunohistochemistry (IHC), and staining

H&E, p53, p21, p-H2AX, and Picro Sirius Red (PSR) staining were performed as previously described^41^. Slides contained 4 µm-thick formalin-fixed paraffin-embedded (FFPE) sections and were heated at 60⁰C for 2 hours before staining was undertaken. Staining was performed in batches on a rolling basis as samples became available for processing and analysis. The exception to this was MDM2 kidney blocks for PSR staining. These were cut and stained in a single run. As much as possible, staining batches included sample tissue from multiple cohorts.

All sections, except for those from the distal organs of female p53 cohort mice (Figure 4 and Figure S5) were scanned at 20x magnification using a Leica Aperio AT2 slide scanner (Leica Microsystems, UK) as previously described^42^. The distal organs of female p53 cohort mice were scanned at 40x magnification using a NanoZoomer-SQ whole slide imaging system (Hamamatsu Photonics, Japan) according to the manufacturer’s recommendations.

### IHC and Staining Quantification

For all liver and kidney p53^+^ cells per tissue area and liver p21^+^ cells per tissue area calculations quantification was conducted using HALO image analysis software (V3.6.4134, Indica Labs) using adaptations of the CytoNuclear module. Kidney p21^+^ cell numbers were obtained from MDM2 cohorts (Figures 2 and S3) using HALO as above. Kidney p21^+^ cells were hand counted in the diet model (Figure 4) as positive cells per 20x field of view. These numbers were averaged across five random fields per sample. This was undertaken due to high background staining observed in HFHSHC samples which confounded the HALO-based approach.

Analysis of PSR and p-H2AX was carried out using the pixel classification tools on QuPath software (v0.6.0)^53^ and reported as percentage of stained area per tissue section. To ensure the robustness of this approach, tissue classification and quantification in QuPath were conducted on a subset of samples independently by two researchers, each obtaining the same trends and similar significance.

Liver steatosis was quantified using the pixel classification tool in QuPath to distinguish macrosteatotic regions, areas of hepatocellular ballooning and microsteatosis, and normal tissue. Steatotic regions obtained from the classifier were pooled to inform the reported overall percentage of steatotic liver tissue per sample.

### Quantitative RT-PCR

Quantitative RT-PCR analysis of liver lysates was undertaken as previously described^42^ using Taqman Fast Advanced Master Mix and Taqman Gene Expression Assays for *Actb* (Mm00607939_s1)*, Cdkn2a* (Mm00494449_m1), and *Gdf15*(Mm00442228_m1).

### ALT assay

Plasma alanine transaminase (ALT) levels were measured from snap frozen mouse plasma samples using the ALT Activity Assay kit (Abcam, Cat# ab105134) according to the manufacturer’s recommendations. Samples were diluted 1:4 in ALT assay buffer and analysed in duplicate per sample.

### Cytokine arrays

Analysis of circulating cytokines present in mouse serum samples was performed using the Proteome Profiler Array Mouse XL Cytokine Array Kit (R&D systems, Biotechne, Cat# ARY028) according to the manufacturer’s recommendations. Array experiments were conducted in two batches, each containing n = 2 biological replicates from each experimental group. The volume of serum analysed was determined by the limiting sample volume within each batch. Membranes were visualised using a ChemiDoc Imaging System (Bio-Rad) and quantification performed using Image Lab (Bio-Rad, Version 6.1.0 build 7). To enable pooling of data, individual spot intensities were normalised to the total signal intensity across all experimental array spots within each batch. Z-score normalised values for all 111 targets per array were then calculated and analysed between groups.

### ELISA measurement of plasma GDF15

Levels of GDF15 were determined from snap frozen mouse plasma samples using the Mouse GDF15 ELISA kit (Proteintech, Cat# KE10082) according to the manufacturer’s recommendations. Samples were diluted 1:6 for the assay and analysed in duplicate per sample. GDF15 concentrations were determined against a standard curve using a four-parameter logistic curve as recommended.

### LiverACE spheroid generation, steatotic induction, and analysis

Human LiverACE multicomponent spheroids comprising differentiated HepaRG-116 cells, monocyte-derived macrophages (MOPs), endothelial cells, and hepatic stellate cells were established for 7 days. Spheroids were then exposed for 72 h to lactate, pyruvate, octanoic acid and ammonium chloride (LPON) to induce steatotic conditions, as previously described^54,55^. Matched spheroids and culture supernatants were collected for proteomic and secretomic analysis. Proteomic analysis was performed by the University of Edinburgh Proteomics and Metabolomics Core Facility using DIA-PASEF mass spectrometry on a timsTOF HT platform, with data processed using Spectronaut/directDIA.

Full details of spheroid composition, culture conditions, sample preparation, mass-spectrometry acquisition and downstream analysis are provided in the Supplementary Methods.

### Analysis of publicly available transcriptomics datasets

Human RNA-seq datasets were accessed through the NCBI Gene Expression Omnibus (GEO) portal using GEO accession numbers: GSE135251 and GSE130970^56,57^. The NCBI GEO2R analysis platform was used to determine gene expression levels. Sample groups were defined based on the information included from each submission. Human proteomics data were accessed via the source data linked to the original publication^58^. Z-score normalised values for human proteomics data were calculated and compared between patients with advanced MASH (F3/F4 fibrosis score) and those with less advanced disease (F0-F2 fibrosis score). Fibrosis scoring was provided in the dataset by the original authors and was based on the Brunt fibrosis classification scheme^59^. In this methodology, fibrosis is graded from F0 (no fibrosis) to F4 (cirrhosis). In the intermediate steps, a score of F1 includes pericellular fibrosis, F2 indicates periportal fibrosis, and F3 includes bridging fibrosis^59^. These staging classifiers were also obtained from the RNA-seq datasets and used to define sample groups.

Further analyses were completed using GraphPad Prism 11 (GraphPad). For human proteomics data, Z-score normalised values were analysed using a two-way ANOVA with the two-stage linear step-up procedure of Benjamini, Krieger and Yekutieli to correct for multiple comparisons by controlling the false discovery rate (FDR) at q < 0.05. RNA-seq datasets were analysed using a one-way ANOVA with Tukey’s multiple comparisons test or an unpaired two-tailed t-test with Welch’s correction according to the number of groups being compared. Data points represent individual patients in these analyses.

### Data plotting and statistical analysis

Data were plotted using GraphPad Prism 11 (GraphPad) and presented as mean ± SD with individual data points. To determine statistical significance, ANOVA analyses or t-tests were performed. When two independent variables were assessed, two-way ANOVA analyses with multiplicity-adjusted p-values and Tukey’s multiple comparisons testing were performed. When one independent variable was examined, either an unpaired two-tailed t-test with Welch’s correction or a one-way ANOVA analysis with multiplicity-adjusted p-values and Tukey’s multiple comparisons testing was performed, as appropriate. For cytokine array analysis, Z-score normalised values were analysed using a two-way ANOVA with the two-stage linear step-up procedure of Benjamini, Krieger and Yekutieli to correct for multiple comparisons by controlling the FDR at q < 0.05. Data points from *in vivo* experiments represent individual mice. Data points from *in vitro* experiments represent individual LiverACE spheroid preparations. Underlying assumptions for statistical tests, including of data normality, were assumed to be met although not explicitly examined.

Figures were prepared using Affinity Designer 2 (Serif).

For further information, see supplementary methods.

## RESULTS

### Loss of MDM2 E3 activity results in progressive liver damage and the accumulation of senescent-like features

MDM2 is a RING finger E3 ubiquitin ligase (E3) that binds to p53 and regulates its activity in partnership with MDMX^60^. As an E3, MDM2 facilitates proteasome-mediated degradation of p53^61^. MDM2 can also bind to nuclear p53 and restrict its access to transcriptional machinery^62,63^. Deletion of the p53 binding domain within MDM2, as in *Mdm2^Ex5/6^*^Δ^ mice, disrupts both regulatory pathways and is embryonically lethal, lethal in adult mice within 4-6 days, and lethal in a liver-specific context (*Mdm2^Ex5/6_liv^*^Δ^) in adult mice after acute deletion within a similar timeframe^32,64–66^. To bypass this issue, we have previously described the creation of an *Mdm2^I438K^* genetic model in which conditional expression of an I438K mutation (I440K in humans) in the MDM2 RING domain blocks E2 binding and disrupts MDM2 E3 activity^51^. *Mdm2^I438K^*mice are E3 activity-deficient but retain p53 binding and non-E3-mediated p53 regulatory capacity. As a result, in adult *Mdm2^I438K^* mice, p53 is stabilised but downstream signalling is restrained, allowing for extended survival compared with *Mdm2^Ex5/6^*^Δ^ mice accompanied by a rapid and potent response to p53 activation after stimuli^51^.

In subsequent efforts to refine the *Mdm2^I438K^* model, we observed allele-specific activity within *Mdm2^I438K^* mice (Figure S1 A-C). In C57BL6/J (N4) *Mdm2^I438K^* mice where the neomycin targeting cassette was excised, converting the published ‘tm1’ allele^51^ into the ‘tm1.1’ variant (Figure S1A), adult whole-body induction of *Mdm2^I438K^* was no longer tolerated (Figure S1 B,C). In contrast with the published outbred tm1 strain, tm1.1 mice reached clinical endpoint within 4-6 days due to progressive weight loss alongside attrition of villi observed in the small intestine and histopathological abnormalities in the spleen (Figure S1 B,C). This suggests exacerbation of a gut-specific toxicity within the model that we reported in tm1 mice^51^ that is no longer resolvable in the tm1.1 strain. Nonetheless, as in tm1 *Mdm2^I438K^* mice, tm1.1 *Mdm2^I438K^*mice exhibited normal histology in other organs, including the pancreas and liver, and evidenced histopathological signs of p53 stabilisation and restrained downstream expression of p21 in the liver (Figure S1C,D)^51^, suggesting that activation of *Mdm2^I438K^* in the liver of tm1.1 mice is better tolerated than liver-specific *Mdm2* deletion.

Although liver-specific abrogation of MDM2-p53 binding in adult *Mdm2^Ex5/6_liv^*^Δ^ (*Mdm2^liv^*^Δ^) male mice is lethal within 4-8 days^32,66^; prior to clinical endpoint, *Mdm2^Ex5/6_liv^*^Δ^ loss is sufficient to induce multiorgan senescence^32^. We also observed evidence of liver damage in adult tm1 *Mdm2^I438K^* mice^51^ but this was tolerated in the model, suggesting that MDM2 E3 activity might be involved in mediating liver toxicity or broader senescence induction. To explore this possibility, we generated cohorts of male and female *Mdm2^I438K^* mice from both tm1 and tm1.1 strains to compare against *Mdm2^liv^*^Δ^ mice. Hepatocyte-specific activation of the *Mdm2^I438K^* allele was achieved via tail vein administration of high titre AAV8-TBG-Cre, as previously described within *Mdm2^liv^*^Δ^ mice^32,52^. Importantly, the resulting *Mdm2^I438K^* mice of both strains were viable and exhibited only mild (<10%) weight loss that was well-tolerated out to a least 19 days, the latest timepoint analysed in this study (Figure S1E).

Consistent with previous reports, we observed substantial IHC staining for p53 and robust expression of p21 in both male and female *Mdm2^liv^*^Δ^ mice compared to AAV8-TBG-Null treated controls (hereafter *Mdm2^liv^*^Δ-WT^ mice) four days after induction (Figure 1A-C)^32^. In addition, male *Mdm2^liv^*^Δ^ mice exhibited increased staining for the fibrosis marker Picro Sirius Red (PSR) and elevated plasma levels of Alanine Transaminase (ALT) activity, indicative of fibrotic liver damage (Figure 1D,E). Female *Mdm2^liv^*^Δ^ mice did not exhibit significant fibrosis or elevated plasma ALT levels compared with *Mdm2^liv^*^Δ-WT^ mice, suggesting potential sex-specific differences in tolerance of hepatic p53 stabilisation and potential differences in survival after induction, although this was not examined in our study (Figure 1D,E). In AAV8-TBG-Cre treated male *Mdm2^I438K^* mice (hereafter *Mdm2^I438K_liv^*mice) we similarly observed abundant IHC staining for p53 compared with minimal staining in AAV8-TBG-Null treated *Mdm2^I438K^* mice (hereafter *Mdm2^WT^)*, as expected (Fig1F,G)^51^. Importantly, p53 was stabilised to the same level in male *Mdm2^I438K_liv^*mice from both tm1.1 and tm1 strains and matched the level of p53 stabilisation observed within *Mdm2^liv^*^Δ^ mice (Figure S2A). We also observed robust stabilisation of p53 in female *Mdm2^I438K_liv^*mice that again aligned with p53 levels in our other cohorts and was absent, as expected, in AAV8-TBG-Null treated female *Mdm2^I438K^* mice (Figure S2B-D).

**Figure 1:**
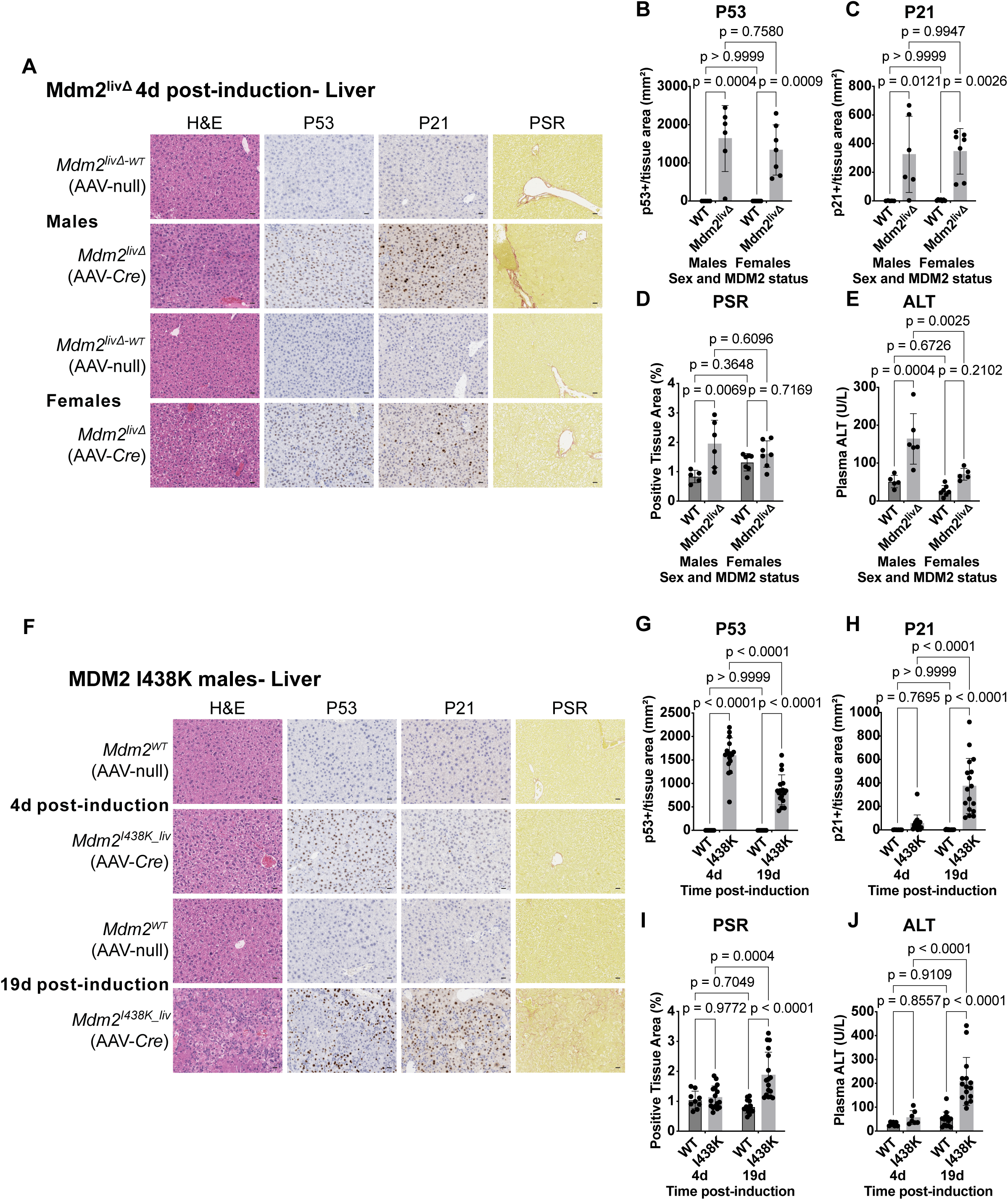
Loss of liver MDM2 E3 activity results in progressive liver damage and the accumulation of senescent-like features in male Mdm2^I438K_liv^ mice. A: Representative liver H&E images and staining for p53, p21, and fibrosis (Picro Sirius Red/PSR) in male and female AAV8-TBG-Cre induced *Mdm2^Ex5/6_liv^*^Δ^ mice (*Mdm2^liv^*^Δ^) and control AAV8-TBG-Null-treated *Mdm2^Ex5/6_liv^*^Δ^ mice (*Mdm2^liv^*^Δ-WT^) mice at 4 days post-induction. N=5 male *Mdm2^liv^*^Δ-WT^ mice, N=6 male *Mdm2^liv^*^Δ^ mice, and N=7 female mice in each group. Scale bars 20 μm. B-D: Quantification of p53 (B) or p21 (C)-positive hepatocytes per tissue area (mm^2^), and PSR positive tissue area (%) (D) in mice from (A). WT: *Mdm2^liv^*^Δ-WT^. N-numbers as in (A). E: Quantification of Alanine Transaminase (ALT) activity (U/L) in plasma samples from mice in (A). N=5 male *Mdm2^liv^*^Δ-WT^ mice, N=6 male *Mdm2^liv^*^Δ^ mice, N=7 female *Mdm2^liv^*^Δ-WT^ mice, and N=5 female *Mdm2^liv^*^Δ^ mice. Samples were not obtained or analysed from N=2 female *Mdm2^liv^*^Δ^ mice. F: Representative liver H&E images and staining for p53, p21, and PSR in male AAV8-TBG-Cre induced *Mdm2^I438K^* mice (*Mdm2^I438K_liv^*) and control AAV8-TBG-Null-treated *Mdm2^I438K^* mice (*Mdm2^WT^)* at 4 or 19 days post-induction. Data are pooled between tm1 and tm1.1 *Mdm2^I438K^*strains in each condition. At 4d, N=9 *Mdm2^WT^* mice and N=16 *Mdm2^I438K_liv^* mice. At 19d, N=14 *Mdm2^WT^* mice and N=17 *Mdm2^I438K_liv^* mice. Scale bars 20 μm. G-I: Quantification of p53 (G) or p21 (H)-positive hepatocytes per tissue area (mm^2^), and PSR positive tissue area (%) (I) in mice from (F). WT: *Mdm2^WT^*. N-numbers as in (F). J. Quantification of Alanine Transaminase (ALT) activity (U/L) in plasma samples from mice in (F). At 4d, N=7 mice per condition. At 19d, N=14 *Mdm2^WT^* mice and N=15 *Mdm2^I438K_liv^* mice. Samples were not obtained or analysed from N=2 *Mdm2^WT^* mice or N=9 *Mdm2^I438K_liv^* mice at 4d or from N=2 *Mdm2^I438K_liv^*mice at 19d. For all graphs, data points are from individual mice and analysed using a two-way ANOVA with Tukey’s multiple comparisons test. Multiplicity-adjusted p-values as shown. Bars show mean ± SD.

In contrast to abundant p53 levels, male *Mdm2^I438K_liv^*mice did not significantly induce p21 expression at 4 days post-induction (Figure 1H). This result is consistent with MDM2^I438K^ maintaining the ability to restrict p53 transcription independent of E3 activity shortly after allele induction^51^. Unexpectedly, however, p21 levels in our female *Mdm2^I438K_liv^* mice were significantly elevated compared to *Mdm2^WT^* controls (Figure S2E), suggesting that MDM2 E3 function may be more important for limiting p53 activity in the female liver. As with p53 staining, neither the level of p21 in *Mdm2^I438K_liv^*mice nor PSR levels differed between tm1.1 and tm1 mice of the same sex (Figure S2F-I), confirming that the two alleles produced similar phenotypes in a hepatocyte-specific context. For this reason, and to maximise the use of the animals in our models, we proceeded to pool observations from tm1.1 and tm1 *Mdm2^I438K^*mice of the same sex together, although we continued to assess both strains for each analysis as described in the methods section.

Liver-specific activation of MDM2^I438K^ was well tolerated in both male and female *Mdm2^I438K_liv^* mice, in contrast to the lethality observed under similar conditions within the *Mdm2^liv^*^Δ^ model^66^. At the same time, while we observed good control over p21 expression in male *Mdm2^I438K_liv^* mice in the short-term, given that this control was not evident in female mice, we questioned whether p53 restriction would persist. To examine this possibility, we analysed liver p53 activity and damage at a later timepoint. At 19 days post-induction, male *Mdm2^I438K_liv^* mice exhibited features consistent with progressive liver damage, including elevated hepatic p21 expression, increased fibrosis, and elevated plasma ALT activity—features that were all absent at 4 days post-induction (Figure 1H-J).

In female *Mdm2^I438K_liv^* mice, we also observed increased fibrosis and persistent elevation of plasma ALT activity (Figure S2J-K), but these occurred alongside sharply reduced p53 stabilisation and decreased (but still elevated) hepatic p21 compared to the 4-day timepoint (Figure S2D-E). These findings suggest different liver damage trajectories in male and female *Mdm2^I438K_liv^* mice following loss of MDM2 E3 activity. This observation was further reinforced through analysis of the DNA damage marker phospho-H2AX (Ser139) (p-H2AX) and expression of *Cdkn2a,* markers of senescence^33^, that were significantly elevated in male *Mdm2^I438K_liv^* liver samples only (Figure S2L-M).

Together, our findings demonstrate that MDM2 E3 activity is dispensable in the short-term for limiting p21 induction, liver damage, and senescent-like features within male *Mdm2^I438K_liv^* mice. However, E3 loss nonetheless facilitated progressive accumulation of liver damage, fibrosis, and senescent features including significant p21 expression, DNA damage, and induced *Cdkn2a* expression in male mice. This was not the case in female *Mdm2^I438K_liv^* mice, where E3 activity appears to be more important for both acute and longer-term control of hepatic p53 activity.

### Loss of MDM2 E3 activity leads to induction of p21 and fibrosis in the kidneys

The progressive liver damage we observed in male *Mdm2^I438K_liv^* mice included features analogous to the senescent state described to occur within male *Mdm2^liv^*^Δ^ mice at 4 days post-liver MDM2 deletion (Figure 1)^32^. One striking consequence of this hepatocellular dysfunction is the reported induction of senescence marked by emergent p21 expression in distal organs including the proximal tubule compartment of the kidneys^32^. To determine if a similar programme was operant within *Mdm2^I438K_liv^* mice, we compared kidney biology across our cohorts from Figure 1. Consistent with published reports, we noted a lack of p53 expression but induction of p21 within the proximal tubule compartment in the cortex of the kidneys in male *Mdm2^liv^*^Δ^ mice (Figure 2A-C)^32^. Neither p53 nor p21 induction was observed in the kidneys of female *Mdm2^liv^*^Δ^ mice, suggesting that transmission of senescent-like features had not occurred (Figure 2A-C). Unlike in the liver, where elevated p21 correlated with fibrosis in male mice, we did not observe evidence of increased PSR staining in the kidneys of either male or female *Mdm2^liv^*^Δ^ mice (Figure 2D).

In male *Mdm2^I438K_liv^* mice, we observed progressive induction of p21 but not p53 within the proximal tubule region of the kidneys that was significant at 19 days post-induction and coincided with increased PSR staining for fibrosis (Figure 2E-H). Female *Mdm2^I438K_liv^* mice, in contrast, did not induce p53, p21, or fibrosis at either timepoint examined (Figure S3A-D). Collectively, our results confirm the induction of systemic p21 that arises downstream of liver senescence in *Mdm2^liv^*^Δ^ mice but provide additional context that this is likely a sex-specific outcome. A similar paradigm occurs in *Mdm2^I438K_liv^* mice, with kidney disruption in male mice taking longer to manifest than within *Mdm2^liv^*^Δ^ mice but including evidence of fibrosis in addition to p21 induction.

**Figure 2:**
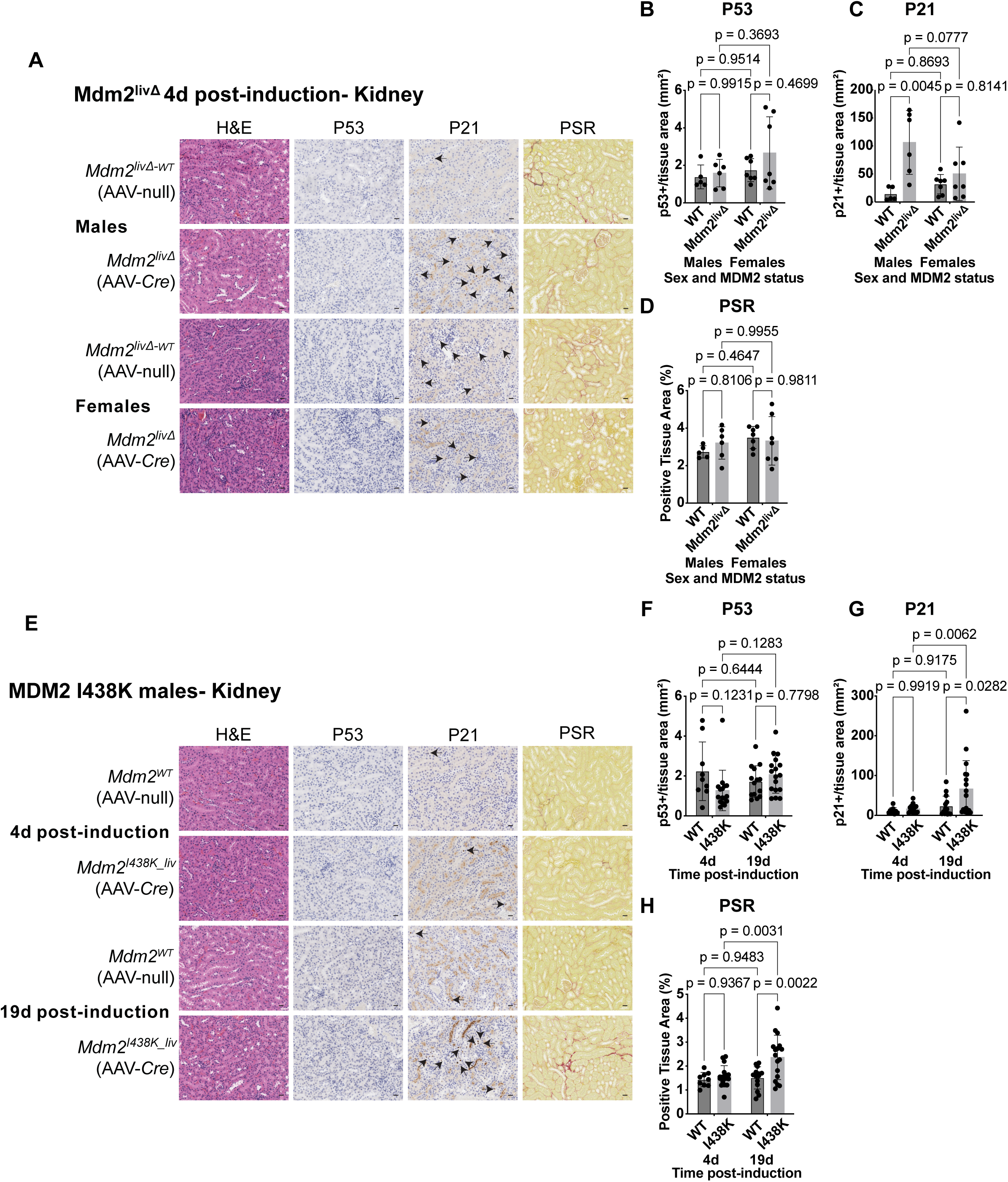
Loss of liver MDM2 E3 activity leads to induction of p21 and fibrosis in the kidneys of male Mdm2^I438K_liv^ mice. A: Representative kidney H&E images and staining for p53, p21, and PSR in male and female *Mdm2^liv^*^Δ^ mice and control *Mdm2^liv^*^Δ-WT^ mice at 4 days post-induction. Images are focused on the cortex and include the proximal tubule compartment. Mice analysed here are the same as presented in Figure 1 for liver analyses. N=5 male *Mdm2^liv^*^Δ-WT^ mice, N=6 male *Mdm2^liv^*^Δ^ mice, and N=7 female mice in each group. Arrows denote p21-positive cells. Scale bars 20 μm. B-D: Quantification of p53 (B) or p21 (C)-positive cells per tissue area (mm^2^), and PSR positive tissue area (%) (D) in mice from (A). WT: *Mdm2^liv^*^Δ-WT^. N-numbers as in (A). E: Representative kidney H&E images and staining for p53, p21, and PSR in male *Mdm2^I438K_liv^* and control *Mdm2^WT^* mice at 4 or 19 days post-induction. Data are pooled between tm1 and tm1.1 *Mdm2^I438K^*strains in each condition. Images are focused on the cortex and include the proximal tubule compartment. Mice analysed here are the same as presented in Figure 1 for liver analyses. At 4d, N=9 *Mdm2^WT^* mice and N=16 *Mdm2^I438K_liv^* mice. At 19d, N=14 *Mdm2^WT^* mice and N=17 *Mdm2^I438K_liv^* mice. Arrows denote p21-positive cells. Scale bars 20 μm. F-H: Quantification of p53 (F) or p21 (G)-positive cells per tissue area (mm^2^), and PSR positive tissue area (%) (H) in mice from (E). WT: *Mdm2^WT^*. N-numbers as in (E). For all graphs, data points are from individual mice and analysed using a two-way ANOVA with Tukey’s multiple comparisons test. Multiplicity-adjusted p-values as shown. Bars show mean ± SD.

### Cholesterol promotes potent hepatocellular p53 activation, MASH, and induction of kidney p21 and fibrosis

Several features of progressive liver damage observed within *Mdm2^I438K_liv^*male mice, including robust expression of p21, persistent DNA damage, and fibrosis are also pathological within human MASH and have been positively linked to hepatocellular senescence^33^. We have previously utilised a whole-body p53 reporter (p53rep) mouse to noninvasively monitor p53 induction in response to an obesogenic high fat high sugar (HFHS) diet leading to MASH over the course of 1 year^42^. However, the slow speed and restricted nature of p53 activity in this model^42^ were not a good match with the rapid and potent p53 signalling observed in *Mdm2^I438K_liv^* mice. Seeking an alternative, we explored additional diet formulations and assessed their impact on p53rep induction *in vivo*.

Cholesterol has been implicated as a substantial driver of hepatocellular damage during liver disease progression^17–22^. In our previous work, we utilised a cholesterol-free HFHS diet which resulted in penetrant but slow-developing liver disease^42^. Within murine models, supraphysiological cholesterol content of 1-2% has been found necessary to accurately model the cholesterol exposure of humans, owing to comparatively poor cholesterol absorption in C57BL/6J mice^22,67^. Consistent with these recommendations and consensus guidelines^68^, we focused on an alternative high fat high sugar high cholesterol diet (HFHSHC) containing 1.27% cholesterol that has previously been reported to cause MASLD to fibrotic MASH transition within 8-16 weeks^69,70^, a significantly enhanced timescale. To compare the effects of these diets on p53 activity, we administered both HFHS and HFHSHC formulations to male and female p53rep+ mice for a period of 75 days (Figure 3A,B & Figure S4A,B). Consistent with our previous work, HFHS-fed p53rep+ mice did not exhibit increased p53rep fluorescence compared with chow-fed p53rep+ mice during this period. HFHSHC-fed male and female mice, in contrast, both exhibited significantly increased p53rep fluorescence relative to baseline by 25 or 50 days of diet, respectively, although p53rep signal intensity was consistently higher in HFHSHC-fed male mice at all time points (Figure 3A,B and Figure S4A,B).

**Figure 3:**
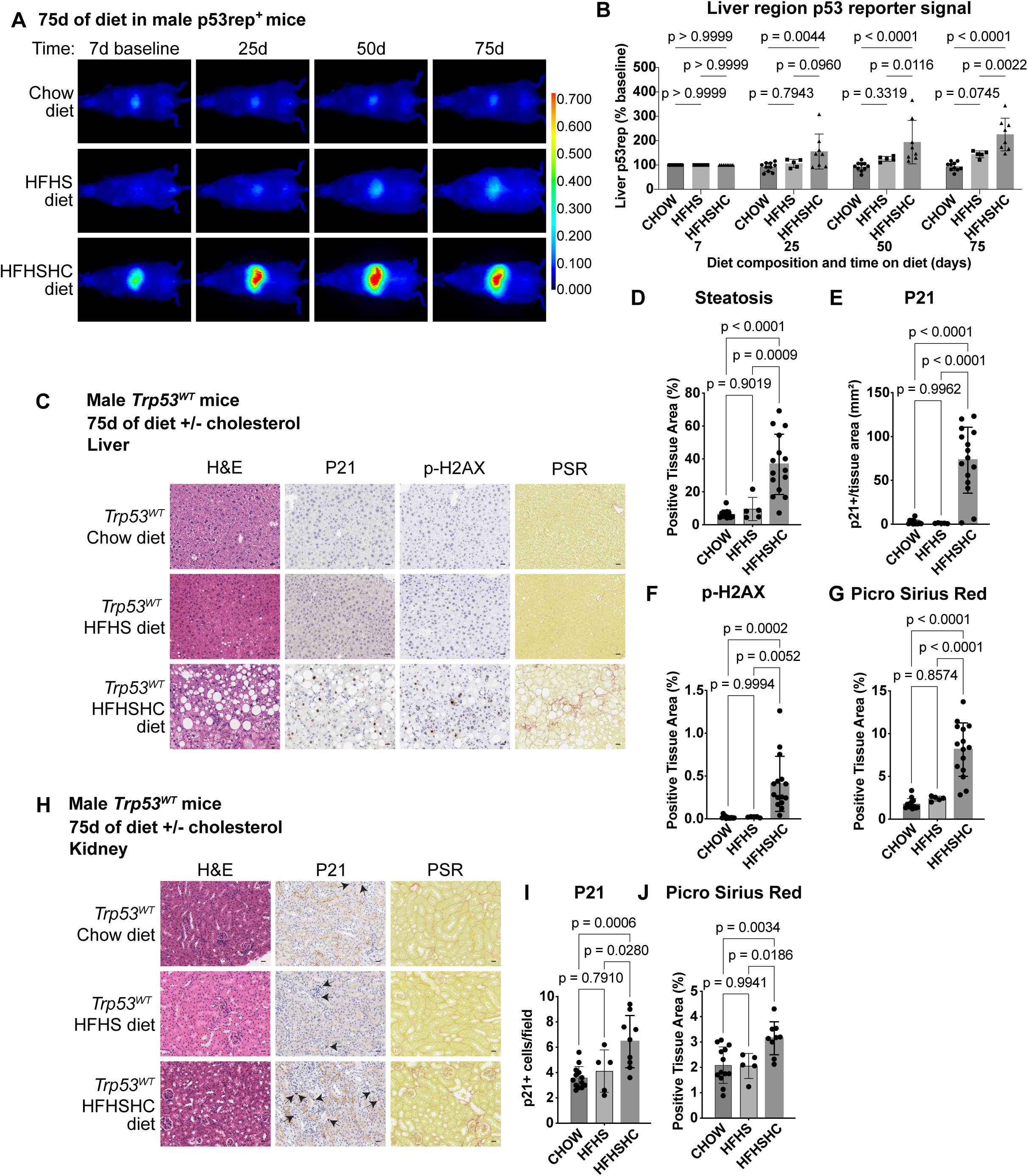
Cholesterol promotes robust liver p53 activation, MASH, and induction of kidney p21 and fibrosis *in vivo*. A/B: Male whole-body p53 reporter (p53rep) mice were given obesogenic high fat high sugar (HFHS) diet or high fat high sugar high cholesterol (HFHSHC) diet at 60-70 days of age or remained on normal chow diet and imaged after the indicated days on diet. Representative images (A) and P53rep signal quantification (B) normalised to the liver signal identified per mouse at the baseline (7d diet) measurement. N=10 chow, N=5 HFHS, and N=8 HFHSHC mice imaged per timepoint. LUT intensity values for images in (A) as shown. Data points are from individual mice and analysed using a two-way ANOVA with Tukey’s multiple comparisons test. Multiplicity-adjusted p-values as shown. Bars show mean ± SD. C. Representative liver H&E images and staining for p21, phospho-H2AX (Ser139) (p-H2AX), and PSR in male *Trp53* WT mice given HFHS, HFHSHC, or control chow diet for 75 days. N=13 chow mice, N=5 HFHS mice, and N=15 HFHSHC mice. *Trp53^WT^*mice comprised a mix of AAV8-TBG-Cre-treated *Trp53* WT mice and uninduced *Trp53^FL/FL^* mice and included both p53rep^+^ and p53rep^-^ mice. Further information in supplemental methods. Scale bars 20 μm. D-G. Quantification of steatosis area (%) in H&E images (D), p21-positive cells per tissue area (mm^2^) (E), or p-H2AX (F) or PSR (G)-positive tissue area (%) in mice from (C). N-numbers as in (C). H-J. Representative kidney H&E images (H) and staining for p21 (I) and PSR (J) in male *Trp53* WT mice given HFHS, HFHSHC, or control chow diet for 75 days. Images are focused on the cortex and include the proximal tubule compartment. N=13 chow mice, N=5 HFHS mice, and N=9 HFHSHC mice. Data are from the cohorts in (C), but kidney was not sampled from all HFHSHC-fed mice in (C). Arrows denote p21-positive cells. Scale bars 20 μm. For graphs in D-J, data points are from individual mice and analysed using a one-way ANOVA with Tukey’s multiple comparisons test. Multiplicity-adjusted p-values as shown. Bars show mean ± SD.

To further verify the role of cholesterol in inducing p53, we developed a reformulated version of our HFHS diet that contained added cholesterol content of 1.27% (HFHS+Chol) but was otherwise as equivalent as possible. This setup isolated the contribution of cholesterol from other factors of the HFHSHC diet formulation. As with HFHSHC-fed mice, male p53rep+ mice on the HFHS+Chol diet exhibited significantly increased liver-region fluorescence compared to HFHS-fed mice after 50 days of diet (Figure S4C,D). *Ex vivo* analysis of tissues from chow, HFHS, HFHSHC, and HFHS+Chol-fed male mice confirmed similar liver specificity and magnitude of induced p53rep signal in HFHSHC and HFHS+Chol-fed mice (Figure S4E,F). Together, these findings suggest that cholesterol significantly enhances induction of liver p53 in the context of a high fat diet, leading to substantial p53 activation within 75 days of diet administration in p53rep+ mice.

In agreement with p53rep findings, histopathological analyses confirmed extensive IHC staining for p21 alongside significant steatosis in male *Trp53* WT HFHSHC-fed mice that was not present in either *Trp53* WT HFHS or chow-fed animals at this time point (Figure 3C-E). Alongside elevated p21 staining, HFHSHC-fed mice also exhibited increased DNA damage and extensive fibrosis (Figure 3F,G) that were absent in chow or HFHS-fed mice. These traits were confirmed in HFHS+Chol-fed *Trp53* WT male mice where p21, DNA damage, and fibrosis were increased compared with HFHS-fed mice, although steatosis was not significantly elevated according to our classifier (Figure S4G-K). Together, these observations identify cholesterol as a significant driver of p53 activity during diet-induced liver disease and connect pathological features between male HFHSHC-fed and Mdm2^I438K_liv^ mice.

Pursuing the links between our diet-induced MASH and *Mdm2^I438K^*models further, we compared kidney histology between *Trp53* WT chow, HFHS, and HFHSHC-fed male mice. As in male *Mdm2^I438K_liv^* mice, both p21 expression and PSR staining for fibrosis were elevated in the proximal tubule region of the cortex in HFHSHC-fed mice compared with either chow-fed or HFHS-fed mice (Figure 3H-J). Fibrosis was also increased in *Trp53* WT HFHS+Chol-fed mice compared to HFHS-fed mice, although kidney p21 was not significantly altered between these two groups (Figure S4L-N).

### Hepatocellular p53 is necessary for p21 induction and fibrosis in the liver and distal organs

Considering the connection between potent p53 activation and liver fibrosis that we observed in both *Mdm2^liv^*^Δ^ and *Mdm2^I438K^* mice (Figure 1), we sought to clarify whether hepatocellular p53 was orchestrating the local and distal fibrosis observed within HFHSHC fed mice. To test this hypothesis, we generated cohorts of male and female *Trp53^FL/FL^* mice where hepatocellular *Trp53* deletion was achieved via administration of high-dose AAV8-TBG-Cre (*Trp53^liv^*^Δ^ mice) in a similar fashion to our *Mdm2* genetic models. Following AAV8-TBG-Cre administration, induced mice were left for 6 days to allow any residual effects of viral induction to subside^52^. They were then fed either HFHSHC diet or remained on chow diet for a period of 75 days and were assessed alongside control p53 WT mice (*Trp53^WT^*) that comprised a mix of AAV8-TBG-Cre-treated *Trp53* WT mice and uninduced *Trp53^FL/FL^*mice.

After 75 days of diet, steatosis was similar in HFHSHC-fed mice independent of liver p53 status (Figure 4A,B). As expected, although p21 was strongly induced in HFHSHC-fed *Trp53^WT^* mice, this induction was absent in *Trp53^liv^*^Δ^ mice (Figure 4C). Similarly, while both sets of HFHSHC-fed male mice exhibited elevated p-H2AX staining for DNA damage, increased PSR staining for fibrosis was liver p53-dependent (Figure 4D,E). In female mice, we also observed p53-independent steatosis alongside liver p53-dependent induction of p21 in HFHSHC-fed mice. However, this was not accompanied by significantly increased fibrosis or DNA damage in *Trp53^WT^* mice, although we did note increased DNA damage in *Trp53 ^liv^*^Δ^ female mice (Figure 4F-I and Figure S5A). Combined, these findings confirm that hepatocellular p53 status plays a role in directing fibrosis within HFHSHC diet-induced MASH. However, as in *Mdm2^I438K_liv^* mice, senescent-like features were more strongly engaged in male than female HFHSHC-fed mice. This conclusion was bolstered by evidence of increased *Cdkn2a* expression within the livers of male *Trp53^WT^* HFHSHC-fed mice compared with *Trp53 ^liv^*^Δ^ males, but no difference between female HFHSHC-fed mice (Figure S5B).

**Figure 4:**
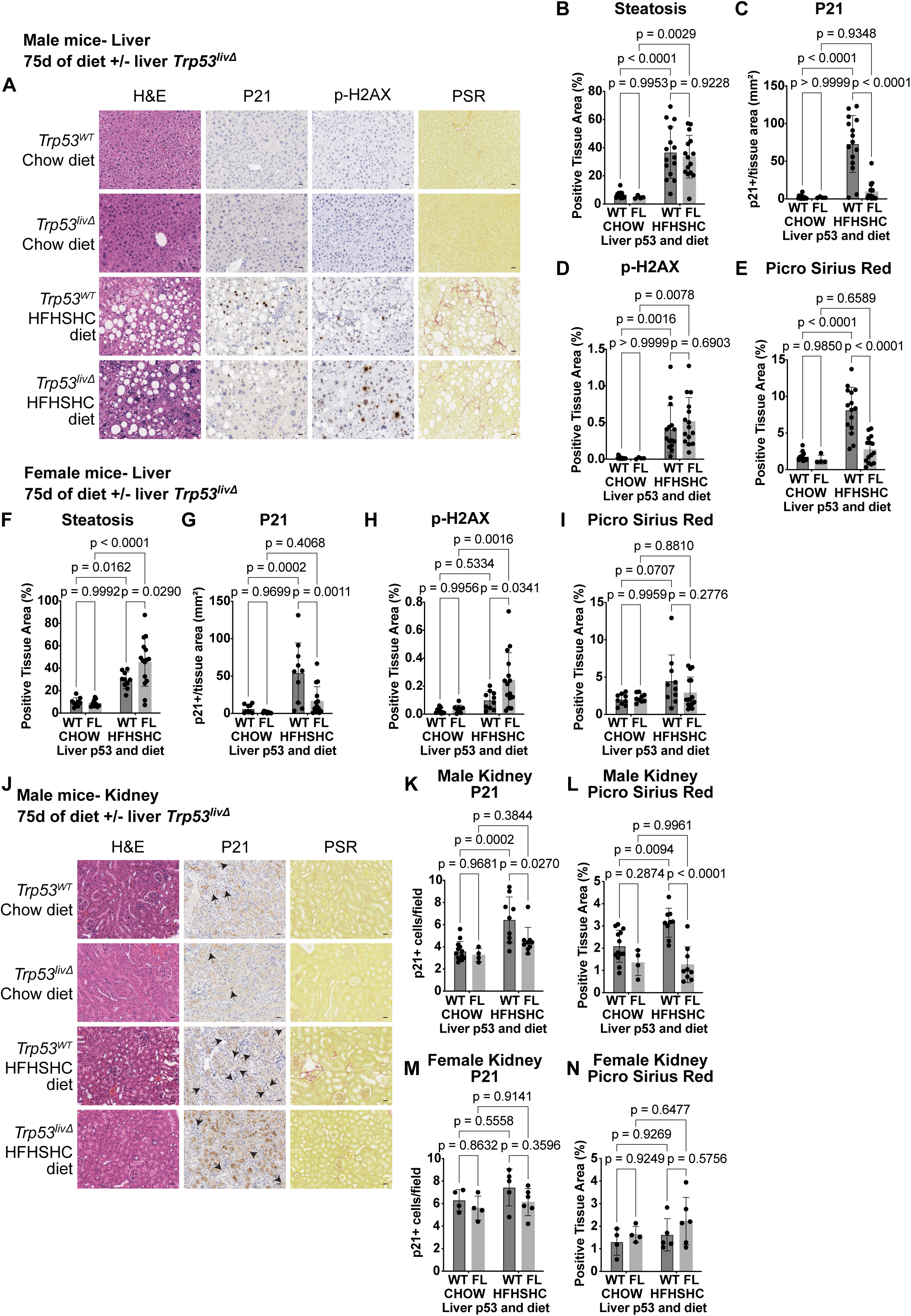
Hepatic p53 enhances liver fibrosis in MASH and is necessary for distal induction of p21 and fibrosis *in vivo*. A: Representative liver H&E images and staining for p21, p-H2AX, and PSR in male AAV8-TBG-Cre induced *Trp53^FL/FL^* (*Trp53^liv^*^Δ^) mice or *Trp53* wildtype (*Trp53^WT^*) mice given HFHSHC or control chow diet for 75 days. N=13 *Trp53^WT^* chow mice, N=4 *Trp53 ^liv^*^Δ^ chow mice, and N=15 HFHSHC mice per genotype. Chow and HFHSHC-fed *Trp53^WT^*mice are the same cohorts as in Figure 3 but different representative images are shown. Scale bars 20 μm. B-E: Quantification of steatosis area (%) in H&E slides (B), p21-positive cells per tissue area (mm^2^) (C), or p-H2AX (D) or PSR (E)-positive tissue area (%) in mice from (A). N-numbers as in (A). WT: *Trp53^WT^*, FL: *Trp53^liv^*^Δ^. Data from WT chow and WT HFHSHC-fed mice are the same as presented in Figure 3 D-G. F-I: Quantification of steatosis area (%) in H&E slides (F), p21-positive cells per tissue area (mm^2^) (G), or p-H2AX (H) or PSR (I)-positive tissue area (%) in female *Trp53^liv^*^Δ^ mice or *Trp53^WT^*mice given HFHSHC or control chow diet for 75 days. N=9 *Trp53^WT^*chow mice, N=9 *Trp53^liv^*^Δ^ chow mice, N=10 *Trp53^WT^* HFHSHC mice and N=15 *Trp53^liv^*^Δ^ HFHSHC mice. J: Representative kidney H&E images and staining for p21 and PSR in male *Trp53^liv^*^Δ^ or *Trp53^WT^* mice given HFHSHC or control chow diet for 75 days as in (A). Images are focused on the cortex and include the proximal tubule compartment. N=13 *Trp53^WT^* chow mice, N=4 *Trp53^liv^*^Δ^ chow mice, N=9 *Trp53^WT^* HFHSHC mice, and N=9 *Trp53^liv^*^Δ^ HFHSHC mice. Kidney was not sampled from all HFHSHC-fed mice in (A). Arrows denote p21-positive cells. Data from WT chow and WT HFHSHC-fed mice are the same as presented in Figure 3C-G but different representative images are shown. Scale bars 20 μm. K/L: Quantification of p21-positive cells per field of view (K) and PSR positive tissue area (%) (L) in mice from (J). N-numbers as in (J). Data from WT chow and HFHSHC-fed mice is the same as presented in Figure 3H-J. M/N: Quantification of p21-positive cells per field of view (M) and PSR positive tissue area (%) (N) in female *Trp53^liv^*^Δ^ or *Trp53^WT^* mice given HFHSHC or control chow diet for 75 days as in (F-I). N=4 chow mice per genotype, N=5 *Trp53^WT^* HFHSHC-fed mice and N=6 *Trp53^liv^*^Δ^ HFHSHC-fed mice. Kidney was not sampled from all cohort mice in (F). For all graphs, data points are from individual mice and analysed using a two-way ANOVA with Tukey’s multiple comparisons test. Multiplicity-adjusted p-values as shown. Bars show mean ± SD.

Fibrosis in human MASH correlates with fibrosis in conditions including CKD, IPF, and CVD^7–12^. Consistent with these aetiological links and our findings in *Mdm2^I438K_liv^*mice, we noted that elevated expression of p21 and increased fibrosis in the proximal tubule region of the kidneys was hepatocellular p53 dependent and only occurred within male mice (Figure 4J-N). We also observed increased PSR staining for fibrosis within the lungs and heart of HFHSHC-fed male *Trp53^WT^* mice that was similarly absent in HFHSHC-fed *Trp53^liv^*^Δ^ male mice and in all female mice, suggesting that liver p53 activity in male mice is required for the induction of kidney p21 and increased fibrosis across multiple organs (Figure 4J-N and Figure S5C-H). Interestingly, not all organs were affected by fibrotic MASH in this way, as we did not observe increased fibrosis in either the spleen or pancreas of HFHSHC-fed *Trp53^WT^* male mice (Figure S5I-J).

Together, these results suggest that hepatocellular p53 contributes to the establishment of fibrotic MASH *in vivo* and is associated with kidney p21 induction and fibrosis across multiple distal organs.

### GDF15 is induced by liver p53 and correlates with markers of fibrotic kidney damage in humans

Considering the local and systemic effects exerted by hepatic p53 activation within *Mdm2^I438K^* mice and our diet-induced MASH model, we hypothesised that secreted factors could provide a unifying molecular link. Utilising cytokine arrays, we identified a collection of 26 factors out of 111 assessed that were differentially present in the serum of male HFHSHC-fed mice compared with chow-fed controls after 75 days of diet (Figures 5A and S6A). Of these, 19/26 were induced, including GDF15, LDL Receptor, and Serpin E1 as the top hits by z-score mean difference (Figure S6A). To refine and validate this list, we assessed the abundance of our 26 murine hits within a published plasma proteomics dataset from human MASH patients reported by Govaere and colleagues in 2023^58^ (Figures 5B and S6B). In the Govaere dataset, levels of 7/25 of our murine hits were also significantly changed in advanced MASH plasma (F3-F4 fibrosis score) compared with less advanced disease (F0-F2 fibrosis score) and 1 murine hit was not present in the dataset^58^. Here, the three top 3 hits by z-score mean difference were: E-Selectin (SELE), TNFRSF11B, and GDF15 (Figures 5B & S6B).

**Figure 5:**
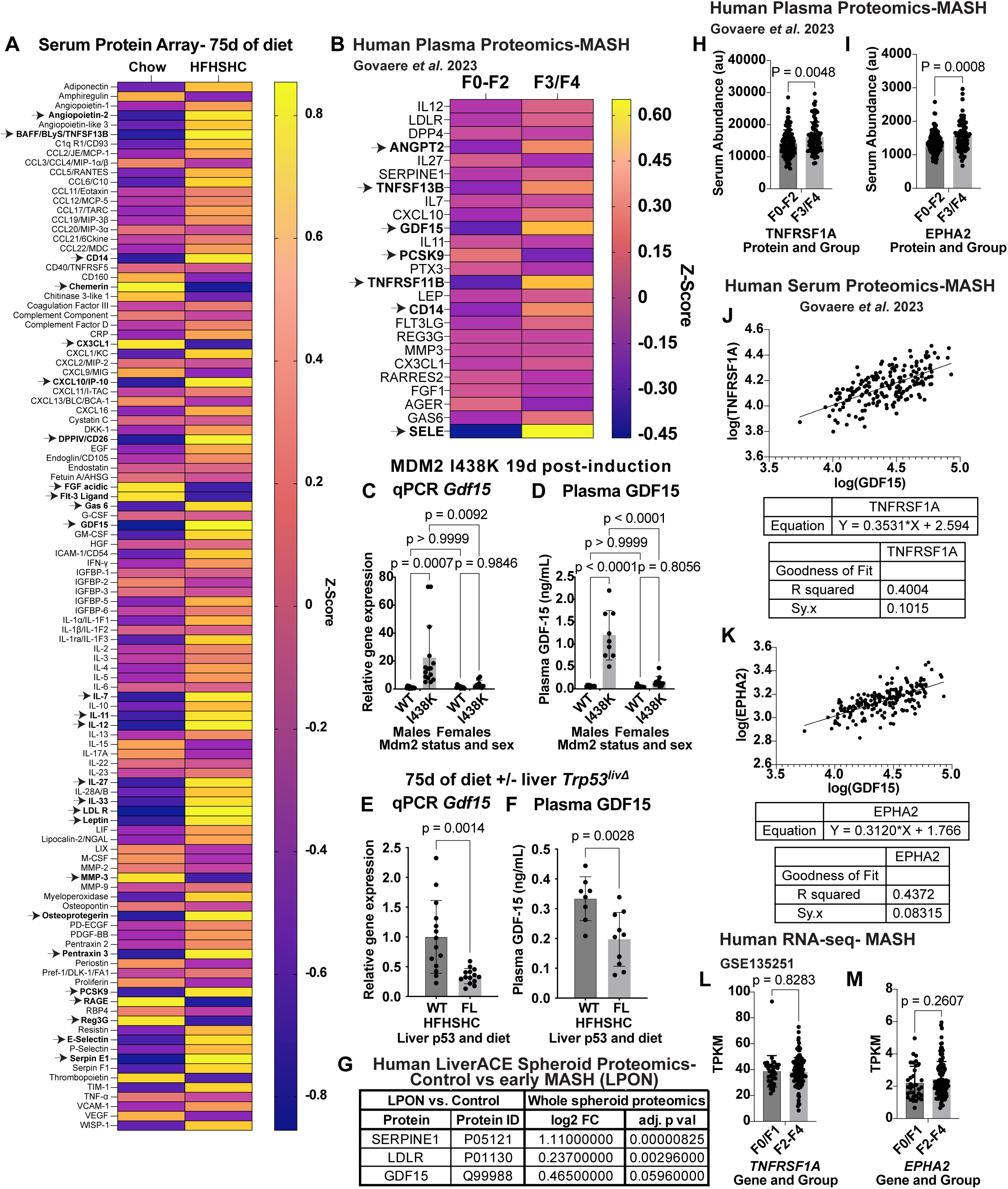
GDF15 is induced by hepatic p53 and correlates with markers of fibrotic kidney damage in humans. A: Proteome Profiler Mouse XL Cytokine Array comparing serum from male mice given HFHSHC or control chow diet for 75 days. Z-score normalised values for all 111 targets per array displayed. N=4 mice per condition. Data analysed using a two-way ANOVA with the two-stage linear step-up procedure of Benjamini, Krieger and Yekutieli to correct for multiple comparisons by controlling the FDR at q < 0.05. Arrows and bolded text identify significant circulating proteins at this threshold. B: Analysis of significant circulating proteins identified in (A) within the published plasma proteomics dataset from human MASH patients reported by Govaere and colleagues in 2023. In the Govaere *et al.* dataset, 25/26 significant circulating proteins from (A) were detected and compared between patients with advanced MASH (F3/F4 fibrosis score) and those with less advanced disease (F0-F2 fibrosis score). Further information on fibrosis score in Materials and Methods. N=112 F0-F2 and N=79 F3/F4 samples. Data presented and analysed as in (A). C: RT-qPCR analysis of liver *Gdf15* expression relative to *Actin* in male and female *Mdm2^I438K_liv^* and *Mdm2^WT^* mice at 19 days post-induction. N=13 *Mdm2^WT^* and N=15 *Mdm2^I438K_liv^* male mice and N=9 female mice per group. Data points are from individual mice and analysed using a two-way ANOVA with Tukey’s multiple comparisons test. Multiplicity-adjusted p-values as shown. Bars show mean ± SD. D: Quantification of plasma GDF15 (ng/mL) by ELISA in male and female *Mdm2^I438K_liv^* and *Mdm2^WT^* mice at 19 days post-induction. N=10 *Mdm2^WT^* and N=9 *Mdm2^I438K_liv^* male mice and N=9 *Mdm2^WT^* and N=11 *Mdm2^I438K_liv^* female mice. Data presented and analysed as in (C). E: RT-qPCR analysis of liver *Gdf15* expression relative to *Actin* in male *Trp53^liv^*^Δ^ or *Trp53^WT^* mice given HFHSHC for 75 days. N=14 mice per condition. F. Quantification of plasma GDF15 (ng/mL) by ELISA in male *Trp53^liv^*^Δ^ or *Trp53^WT^* mice given HFHSHC for 75 days. N=8 *Trp53^WT^*mice and N=10 *Trp53 ^liv^*^Δ^ mice. For graphs in D-F, data points are from individual mice and analysed using an unpaired two-tailed T-test with Welch’s correction. P-value as shown. Bars show mean ± SD. G. Whole spheroid proteomic mass spectrometry in LiverACE spheroids cultured in baseline medium (control) or in LPON medium for 72 h to induce steatosis. The top 3 (out of N=7 detected) proteins identified in (A) are presented in rank order based on adjusted p-value. N=10 control and N=11 LPON samples. Benjamini–Hochberg-adjusted p-value (adj. p val) and log2 fold change (log2 FC) per protein as shown. For significance, differential proteins were required to meet both the Benjamini–Hochberg-adjusted p<0.05 and |log2FC|≥0.25 criteria. Further information available in Supplementary Methods. H/I: Analysis of plasma TNFRSF1A (H) and EPHA2 (I) between patients with advanced MASH (F3/F4 fibrosis score) and those with less advanced disease (F0-F2 fibrosis score) within the Govaere et al. 2023 dataset in (B). N=112 F0-F2 and N=79 F3/F4 samples. Data points are from individual patients and analysed using an unpaired two-tailed T-test with Welch’s correction. P-value as shown. Bars show mean ± SD. J/K: Linear regression analysis comparing log transformed plasma levels of GDF15 against log transformed levels of TNFRSF1A (J) or EPHA2 (K) including patients with advanced MASH (F3/F4 fibrosis score) and those with less advanced disease (F0-F2 fibrosis score) as in (G/H). N=191, including N=112 F0-F2 and N=79 F3/F4 samples. Data points are from individual patients and analysed using a simple linear regression with derived equation and goodness of fit parameters as shown. L/M: Analysis of *TNFRSF1A* (L) and *EPHA2* (M) liver gene expression in the public transcriptomics dataset GSE135251 comparing MASH patients of low fibrosis score (F0/F1) and MASH patients with high fibrosis score (F2-F4). N=34 F0/F1 and N=121 F2-F4 samples. Data presented in transcripts per kilobase million (TPKM). Data points are from individual patients and analysed using an unpaired two-tailed T-test with Welch’s correction. P-value as shown. Bars show mean ± SD.

Based on overlap, we focused our attention on GDF15 as a potential connection across our models. GDF15 is a TGF-β superfamily member^71^ and established p53 target^72^ that has also been implicated as a mediator of MASH^56,73,74^ and fibrosis in multiple organs^75–78^. *In vivo*, we observed increased expression of *Gdf15* by qPCR in liver lysates from male but not female *Mdm2^I438K_liv^* mice at 19 days post-induction that coincided with significantly increased circulating plasma levels of GDF15 in the males at this time point (Figure 5C,D). Reciprocally, we observed decreased liver *Gdf15* expression by qPCR and reduced circulating GDF15 in the plasma of HFHSHC-fed male *Trp53^liv^*^Δ^ mice compared with WT animals after 75 days of diet (Figure 5E,F).

For further insight into the potential sequencing of hepatic GDF15, we turned to *in vitro* modelling utilising a multicomponent liver spheroid model, LiverACE. The LiverACE model includes five principal cell types of the human hepatic acinus (hepatocytes, cholangiocytes, macrophages, endothelial cells, and stellate cells) cultured together at human-relevant proportions within an *in vitro* 3D spheroid. In this context, we compared spheroids treated for 72 h with lactate, pyruvate, octanoic acid and ammonium chloride (LPON)-enriched medium to induce steatosis and model early MASH^54,55^ compared with control-treated LiverACE samples (Figures 5G and S6C). When assessing all 26 plasma- array targets in LPON spheroids, seven were detected by whole-spheroid LiverACE proteomics, including GDF15. Among these, Serpin E1, LDL Receptor and GDF15 ranked highest by Benjamini–Hochberg-adjusted p value but did not all meet our defined differential protein criteria—a Benjamini-Hochberg-adjusted p<0.05 together with |log2FC|≥0.25. Serpin E1 met both criteria (adjusted p=8.3×10⁻⁶; log2FC=1.110); LDL Receptor met the adjusted-p criterion but fell below the effect-size cutoff (adjusted p=0.00296; log2FC=0.237); and GDF15 had a positive effect estimate above the effect- size cutoff but did not quite meet the adjusted-p criterion (adjusted p=0.0596; log2FC=0.465) (Figures 5G and S6C). GDF15 was also detected in culture medium but was not significantly increased following LPON treatment (adjusted p=0.733; Figure S6C), suggesting that local expression of GDF15 in early MASH may be proximal to significant release of this factor later in the progression of the disease.

Across our experimental systems, significant hepatocellular p53 pathway activity was sufficient to promote kidney dysfunction as assessed by induced p21 and fibrosis in the proximal tubule compartment (Figures 2-4). Unfortunately, patient-level data on kidney function was not available within the published Govaere *et al*. plasma proteomics dataset. Seeking a surrogate marker for kidney damage, we examined the plasma levels of TNFRSF1A/TNFR1 and EPHA2, two markers that have been implicated in both chronic kidney disease and acute kidney damage^79–82^. In the MASH dataset, both TNFRSF1A and EPHA2 were significantly increased in the plasma from advanced MASH patients and both positively correlated with plasma GDF15 levels across MASH disease state (Figure 5H-K). Importantly, while *GDF15* expression was induced with increasing fibrosis score alongside *p21* and *CDKN2A* in multiple published human MASH liver transcriptomic datasets^56,57^, we found that *TNFRSF1A* and *EPHA2* were not induced in the liver, consistent with an extrahepatic source for their increased plasma levels, as proposed (Figures 5L,M & S6D-K).

Collectively, these findings identify GDF15 as a p53-dependent hepatic and circulating factor *in vivo*, with a concordant but non-FDR-significant increase in whole-spheroid LiverACE proteomics and increased abundance in advanced human MASH, where circulating GDF15 correlates with markers linked to kidney injury.

## DISCUSSION

P53-mediated tumour suppression is a powerful bulwark against transformation, including within the liver^83^. Disruption of the p53 pathway is associated with worse patient outcomes in HCC and promotes resistance to immune checkpoint-based therapy^84,85^. Roles for p53 within the context of liver damage and repair are significantly more mixed. The p53 pathway has been described to protect the liver during injury by limiting inflammation, reducing fibrosis, enhancing protection against reactive oxygen species (ROS)-related damage, and protecting from diet-induced apoptosis^36–41^. On the other hand, increased p53 pathway activity is correlated with advanced MASH, liver fibrosis, and an increased risk of multiorgan failure in patients^32,34,35^. These paradoxical outcomes likely reflect context-specific balancing of p53 activity^43–45^. In such a model, p53-mediated protection is achieved by modest activation of p53 target genes and without durable p53 stabilisation^46–48^. Past a threshold, however, chronic elevation of p53 becomes a potent driver of senescence and cell death that can be detrimental to liver function^32,34,35^.

Our work provides insight into these disparate outcomes. When p53 activation is sufficiently restrained, as in the acute response following MDM2 E3 loss, we have found that liver function is protected and hepatocellular damage is limited. This finding is reminiscent of the protective p53 response observed in regeneration^41,86^, against alcoholic fatty liver disease^87^, and in response to an obesogenic high fat high sugar diet that lacks cholesterol, as we have previously reported^42^ and confirmed here. Sustained activation of the p53 pathway, in contrast, such as following loss of MDM2-p53 binding, progressively after MDM2 E3 activity is compromised, or in response to a high fat high sugar diet with cholesterol is a common driver of fibrotic liver damage alongside features of hepatocellular senescence. Here, our findings are consistent with reports linking elevated p53 activity with hepatocellular senescence in MASH and an established connection between liver- derived GDF15 and MASLD to MASH progression^56,73,74^. These findings should also be considered within the context-dependent biology of GDF15. In acute inflammatory settings, GDF15 has been reported to promote tissue tolerance and systemic metabolic adaptation^88^; whereas in chronic metabolic disease, sustained elevation may reflect ongoing tissue stress and compensatory responses^89,90^ and not necessarily a pro-fibrotic role.

By dissecting sex-specific responses to both E3 loss and a MASH diet, our findings add nuance to the picture of p53-mediated hepatocellular senescence and the MDM2-p53 regulatory relationship. Our results confirm that disruption of the MDM2-p53 binding interaction in the liver promotes hepatocellular senescence, as reported ^32^, but clarify that this occurs within male but not female mice. We observe a similar sex-dependent difference in the response to a MASH diet wherein female mice are again resistant to downstream deleterious consequences of p53 activation. These results are consistent with broader evidence on sex differences in senescence and may reinforce the concept of sex disparity in p53 regulation, tolerance of p53 activity, or hormone-associated differences in p53 signalling between males and females^91–93^. Future work in this area could provide insight into broader sex-related differences in MASH and MASH-HCC aetiology between male and female patients, an area where pre-clinical models have important utility^94^.

Patients with increased multiorgan fibrosis have a significantly increased risk of mortality than those with limited fibrosis^31^. Even so, molecular understanding of shared mechanisms promoting systemic fibrosis remains incomplete. We have identified fibrosis in distal organs as a conserved pathological consequence of chronically elevated hepatic p53 signalling in our *in vivo* models. These findings are consistent with the interrelationship observed in patients between fibrotic MASH and CKD, CVD, and IPF-related fibrosis and are compatible with contributions from a common and potentially liver-centric pathologic driver ^7–12^. To this end, our work highlights a correspondence between circulating hepatocellular p53-dependent SASP factors including GDF15 and broader multiorgan induction of p21 and increased fibrosis *in vivo*. Using published datasets, we note that GDF15 is elevated in advanced human MASH, consistent with published reports^56,73,74^, and highlight a positive correlation between GDF15 and circulating TNFRSF1 and EPHA2, markers of kidney damage and systemic fibrosis, within MASH patients^79–82^.

Together, these results reinforce a putative molecular link between cholesterol, fibrotic MASH, and distal fibrosis that could potentially be used to identify or monitor patients at- risk of developing multiorgan comorbidities. Further work in this area will be important for clarifying the main contributing factors to multiorgan fibrosis in MASH and determining whether they are therapeutically actionable. Of note, GDF15 and TNFSF13B, another significantly enriched circulating protein from our analyses, have also been highlighted as two of fourteen biomarkers in a powerful predictive panel for lung adenocarcinoma risk that also correlates with increased risk of inflammatory respiratory conditions in humans^95^. Further exploration of this connection as a potential multi-disease risk stratification metric is warranted. As are studies evaluating the efficacy of targeting hepatic p53 alongside diet or therapeutic interventions. Such an approach could provide new treatment approaches for multiorgan fibrosis and MASH.

## ACKNOWLEDGEMENTS

We thank the Core Facilities and Advanced Technologies at the CRUK Scotland Institute (ROR 03pv69j64), and particularly the Biological Services facilities staff (RRID:SCR_028654), the Beatson Advanced Imaging Resource (RRID:SCR_023875), the histology team (RRID:SCR_027306), and the research integrity team (RRID:SCR_028431). We also thank Catherine Winchester (CRUK Scotland Institute) for insightful comments on the manuscript and Mark Thomas Shaw Williams (Glasgow Caledonian University) for constructive feedback throughout the project. The graphical abstract was created in BioRender (Wittke, C. (2026)); https://BioRender.com/23dbw3u.

## AUTHOR CONTRIBUTIONS

CRediT (Contributor Roles Taxonomy) author statement:

**CIW** and **DMW:** Methodology, Investigation, Formal Analysis, Writing-Original Draft Preparation and Review & Editing; **LB, AML, CJC:** Investigation, Formal Analysis, Writing-Review & Editing; **CM, NC, AL, DA, AY, CN:** Investigation, Writing-Review & Editing; **DS:** Resources, Writing-Review & Editing; **LJN**: Resources, Supervision, Writing-Review & Editing; **TGB, KHV:** Conceptualization, Resources, Supervision, Writing-Review & Editing; **KB:** Conceptualization, Supervision, Funding acquisition, Writing-Review & Editing; **JC and TJH:** Conceptualization, Investigation, Formal Analysis, Writing-Original Draft Preparation and Review & Editing, Supervision, Project Administration, Funding Acquisition.

## FUNDING

This work was supported by the United Kingdom Medical Research Council [MR/X018512/1] (awarded to JC and TJH) and the Academy of Medical Sciences [AMS] Springboard scheme [SBF008\1034] (awarded to TJH and funded CIW), which is joint funded by the AMS, the Wellcome Trust, the Government Department of Business, Energy and Industrial Strategy (BEIS), the British Heart Foundation and Diabetes UK. Additional support was provided by the CRUK Scotland Institute which receives its core funding from Cancer Research UK [CRUK grant A31287]. LJN and AML are supported by UKRI Horizon Grant Halt-RONIN 10067052. CJC receives support from CRUK (PRCBTP-May24/100005). TGB was funded by CRUK Accelerator Award (HUNTER: A26813), Wellcome Trust (grant number WT107492Z), and CRUK core funding to the CRUK Scotland Institute (A17196 and A31287). KHV is funded by CRUK grant C596/A26855, ERC-2020-ADG PancObese 10102064 and is supported by the Francis Crick Institute, which receives its core funding from CRUK (CC2073), the UK Medical Research Council (CC2073) and the Wellcome Trust (CC2073). KB receives CRUK Core funding (A29799) which supports KB, AL, and DA. DMW is funded by the Medical Research Council (MC_PC_20142) awarded to KB.

## DECLARATION OF INTERESTS

KHV is on the board of directors and shareholder of Bristol Myers Squibb and on the science advisory board (with stock options) of RAZE Therapeutics, Volastra Pharmaceuticals and Kovina Therapeutics. She is a co-founder, consultant and shareholder of Faeth Therapeutics, a company exploring the impact of non-essential amino acid restriction on therapy in patients. She is also on the advisory board of Cell, Cancer Cell, Molecular Cell, Cell Metabolism and Cell Press Blue. She has been in receipt of research funding from AstraZeneca and contributed to CRUK Cancer Research Horizons’ filing of patent application WO/2017/144877. TGB has received research funding from Iterion Therapeutics, Diogenx and Trogenix. The other authors declare no competing interests.

## DATA AND MATERIALS AVAILABILITY

All data needed to evaluate the conclusions in the paper are present either within the paper itself or the included Supplementary Materials. Underlying raw data from this study are available from the corresponding author upon reasonable request. No new code was generated in this study. All mass spectrometric data concerning human LiverACE spheroids will be made available publicly upon publication as part of Horizon-Europe Grant Agreement ID: 101095679.

---

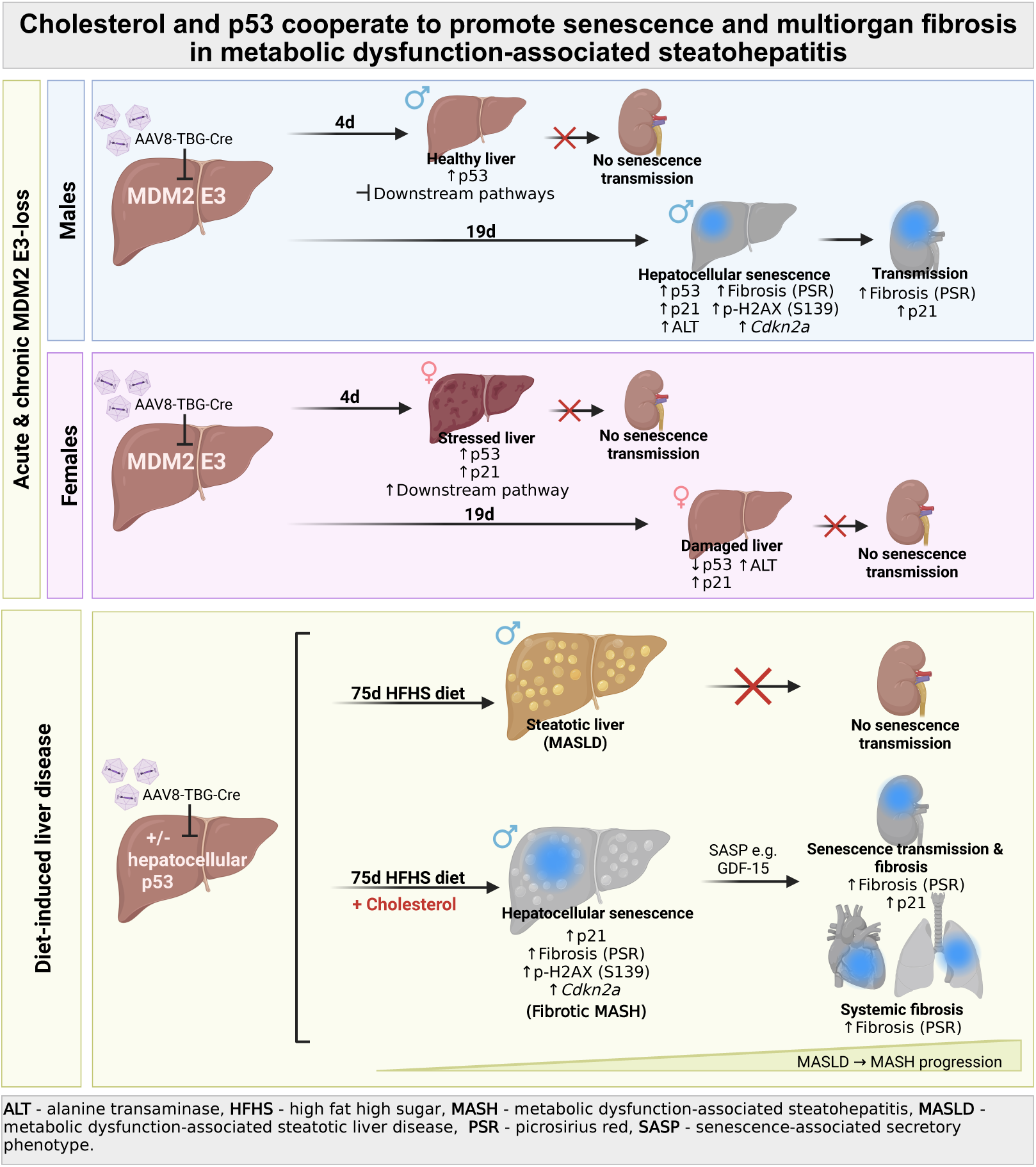

