## Supplementary Materials for "Cholesterol and p53 promote senescence and systemic fibrosis in metabolic dysfunction-associated steatohepatitis"

Running title: Cholesterol and p53 promote senescence and multiorgan fibrosis in MASH

Keywords: p53, MDM2, fibrosis, MASH, cholesterol, GDF15, senescence

#### TABLE OF CONTENTS


1. **Supplementary Methods**
2. **Figure S1:** Whole-body induction of *Mdm2*$^{I438K}$ in adult mice has divergent strain and allele-specific consequences
3. **Figure S2:** The hepatocellular response to loss of MDM2 E3 activity is similar across Mdm2$^{I438K}$ alleles but differs between male and female Mdm2$^{I438K_liv}$ mice
4. **Figure S3:** Loss of liver MDM2 E3 activity does not alter p21 or fibrosis in the kidneys of female Mdm2$^{I438K_liv}$ mice
5. **Figure S4:** Cholesterol promotes robust liver p53 activation and MASH *in vivo*
6. **Figure S5:** Hepatic p53 enhances liver fibrosis in MASH and is necessary for distal induction of p21 and fibrosis *in vivo*
7. **Figure S6:** GDF15 is induced by hepatic p53 and correlates with markers of fibrotic kidney damage in humans
8. **Supplementary Materials References**

#### SUPPLEMENTARY METHODS

##### Additional information relating to *in vivo* experiments

Stock animals were housed in groups of 3-5 as much as possible in individually vented cages and were provided with normal chow diet and water *ad libitum*. Cohort mice were housed in groups of 2-5 mice as much as possible in open-topped cages and provided with formulated diet or normal chow diet and water ad libitum. Non-aversive handling techniques were utilised throughout each experiment. Mice were genotyped by Transnetyx (Cordova, TN).

The strain background of cohort p53 mice in figures 3-5 was analysed through MiniMUGA Background Analysis (v2.3.1, Transnetyx), confirming the inbred status of the colony (98-100% inbred) and its C57BL/6J background (~98% of the genome) excepting portions of the X chromosome and chromosome 11 where the expected 129P2/OlaHsd PG13-iRFP and $p53^{FL/FL}$ constructs, respectively, were targeted.

Tm1 $Mdm2^{I438K}$ mice were confirmed as outbred with a likely mix of C57BL/6J, C57BL/6NTac, and 129P2/OlaHsd background strains and approximately 65-75% C57BL/6J contribution according to MiniMUGA Background Analysis. Tm1.1 $Mdm2^{I438K}$ mice were backcrossed to C57BL6/J (N4) and assessed as close to inbred (95-97% inbred) with a mix of C57BL/6J SNPs (91-94%) and 129P2/OlaHsd SNPs (4-7%), including expected 129P2/OlaHsd contributions from the $Mdm2^{I438K}$ and *Rosa*-Cre$^{ER}$ (Gt(ROSA)26Sor$^{tm2(cre/ERT2)Brn}$) constructs on chromosomes 10 and 6, respectively.

Within experimental cohorts, mice were age and littermate matched as much as possible. MDM2 cohorts, including $Mdm2^{Ex5/6\Delta}$ and $Mdm2^{I438K}$ strains, were started on experimental procedures at approximately 10 weeks of age, with a range of 69 - 82 days of age. Cohort p53 mice, including $Trp53^{liv\Delta}$ and $Trp53^{WT/WT}$ mice started diet interventions between approximately 9 and 11 weeks of age, with the majority starting between 60 and 70 days of age. Most $Trp53^{WT}$ cohorts comprised a mix of uninduced *Trp53* WT and uninduced $Trp53^{FL/FL}$ mice. The exceptions to this were female $Trp53^{WT}$ chow-fed mice and male $Trp53^{WT}$ HFHSHC-fed mice where approximately half of each cohort included $Trp53^{WT}$ mice treated with AAV8-TBG-Cre. These did not differ from uninduced WT cohort mice, as expected, considering the limited and short-term effects of AAV8-TBG-Cre treatment on liver biology previously described[1]. In addition, the majority of $Trp53^{WT}$ and $Trp53^{liv\Delta}$ cohorts included a mix of p53rep$^+$ and p53rep$^-$ animals. This did not affect the response to diet. All p53rep$^+$ males were hemizygous for the allele while p53rep$^+$ females comprised a mix of homozygous and heterozygous mice.

For whole-body induction of $Mdm2^{I438K}$ in Figure S1, *Rosa*-Cre$^{ER+}$ mice were treated via subcutaneous (SC) injection with tamoxifen daily for four days (3 mg of tamoxifen on day 1 or 2 mg of tamoxifen on days 2-4) as previously described[2].

For liver-specific Cre-mediated allele induction, mice were treated with AAV8-TBG-Cre via tail vein injection at a dose of $2 \times 10^{11}$ genomic copies (GC) per mL in a final volume of 100 µL sterile PBS. MDM2 cohorts all received either AAV8-TBG-Cre or AAV8-TBG-Null control treatment of similar titre. Within p53 cohort mice, all $Trp53^{liv\Delta}$ mice received AAV8-TBG-Cre treatment.

For the determination of sample sizes, findings from preliminary small cohorts were used to gauge relative effect size. Power calculations were then performed with G*Power (version 3.1.9.6)[3], using an ANOVA test to compute sample size. Based on an anticipated large effect size (f=0.75 by Cohen's definition), power of 0.90, and, in the case of the 75-day diet cohorts in Fig. 4, for example, four total groups per sex, this analysis suggests a total N of at least 32, equating to 8 mice per experimental group. Similar power calculations were performed for our other *in vivo* experiments and guided our cohort generation strategies.

**Additional information relating to diet experiments**

The HFHS diet (TestDiet 58R3) was cholesterol-free and contained 59.4% energy from fat, 14.9% energy from protein, and 25.7% energy from carbohydrates. The HFHSHC diet (TestDiet 5ZSF) contained 1.27% cholesterol, 52.4% energy from fat, 10% energy from protein, and 37.5% energy from carbohydrates. The HFHS+Chol diet (TestDiet 5ZY4) was modified by TestDiet from 58R3 and contained 1.25% cholesterol, 60.2% energy from fat, 14.7% energy from protein, and 25% energy from carbohydrates. The control normal mouse chow (DS801752G10R, Special Diet Services) was cholesterol-free and contained 9% energy from fat, 21% energy from protein, and 70% energy from carbohydrates. This is an alfalfa-free diet and was previously chosen as being suitable for imaging studies to ensure no/minimal background fluorescence in the near infrared region of the spectrum[4]. The HFHS, HFHSHC, and HFHS+Chol diets were similarly refined and alfalfa-free.

**Additional information relating to Histology, Immunohistochemistry (IHC), and staining**

All haematoxylin and eosin (H&E), IHC, and picro sirius red (PSR) staining for MDM2 cohort mice was performed using formalin fixed paraffin embedded (FFPE) multiblocks. These multiblocks typically contained tissue from 3-4 mice per block and were constructed by separately processing liver or kidney tissue from different mice using a standard histological processing schedule. The individual tissue pieces were then arranged and embedded in histological wax (GCA-08000-25A, CellPath) in a known pre-determined order allowing the location of each piece of tissue to be specifically known within the tissue multiblock and on the resulting glass section. The tissue multiblock approach was also utilised for liver sections from p53 cohort mice. Analyses of distal organs were performed on tissue samples individually blocked for each mouse, with one block containing pancreas, spleen, kidney, and an additional liver sample and the second block containing the heart and lungs.

**Additional information relating to IHC antibodies and staining information**

| **Name** | **Supplier** | **Catalogue number** | **Clone number** |
|---|---|---|---|
| P53 | Leica | NCL-L-p53-CM5p | CM5 |
| P21 | Abcam | ab107099 | HUGO291 |
| Phospho-histone H2A.X (Ser139) | Cell Signaling Technology | 9718 | 20E3 |

| **Name** | **Autostainer** | **Retrieval** | **Dilution** |
|---|---|---|---|
| P53 | Agilent Autostainer Link48 | TRS High | 1/750 |
| P21 | Leica Bond Rx | ER2 20 mins | 1/250 |
| Phospho-histone H2A.X (Ser139) | Leica Bond Rx | ER2 10 mins | 1/120 |

**Additional information relating to LiverACE spheroid generation and steatotic induction**

Human hepatic acinar (LiverACE) spheroids were seeded at a density of 2,000 cells per spheroid in individual wells of a Nunclon Sphera 96-well U-shaped microplate (174925, Thermo Fisher Scientific). Each spheroid contained an initial seeded composition of 70% cryopreserved differentiated HepaRG-116 cells (HPR116080, Biopredic International), 5% cryopreserved primary monocyte-derived macrophages (MOPs; MOP013032, Biopredic International), 15% human umbilical vein endothelial cells (HUVECs) (HUV01028, Biopredic International), and 10% frozen stellate cells (STL101028, Biopredic International).

Spheroids were seeded and cultured for 3 days in a 1:1 mixture of William's E Medium with GlutaMAX™ (32551020, Thermo Fisher Scientific) + HepaRG Thawing/Plating/General Purpose Medium Supplement with antibiotics (ADD670C, Biopredic International) and EBM-2 Basal Medium (CC-3156, Lonza) + EGM-2 SingleQuots Supplements (CC-4176, Lonza). After 3 days, the culture media was replaced with a 1:1 mixture of William's E Medium with GlutaMAX + HepaRG Maintenance and Metabolism Medium (MMM) Supplement with Antibiotics (ADD620C, Biopredic International) and EBM-2 Basal Medium + EGM-2 SingleQuots Supplements. After 7 days, spheroids were cultured in MMM:EBM-2 alone (Control) or MMM:EBM-2 medium supplemented with 10 mM sodium L-lactate (L7022, Sigma-Aldrich), 1 mM sodium pyruvate (P5280, Sigma-Aldrich), 2 mM octanoic acid (C2875, Sigma-Aldrich), and 4 mM ammonium chloride (A4514, Sigma-Aldrich) (LPON) for 72 h to induce steatosis, as previously described[5,6]. Immediately following this, the culture medium supernatants per spheroid and each matched individual spheroid were collected in separate Protein LoBind tubes (EP0030108442, Eppendorf), snap frozen, and stored at -70°C.

**Additional information relating to whole-spheroid proteomics and culture-medium secretomics by mass spectrometry in human hepatic acinar (LiverACE) spheroids**

Proteomic sample preparation for individual LiverACE spheroids and matched cell media supernatants was performed by the University of Edinburgh’s Proteomics and Metabolomics Core Facility using a surfactant-assisted one-pot (SOP) workflow adapted from Tsai *et al.*[7]. Frozen spheroids were washed with phosphate-buffered saline (PBS) and pelleted by gentle centrifugation and cell lysis was achieved by the addition of 0.2% (w/v) n-dodecyl-β-D-maltoside (DDM) (D4641, Sigma-Aldrich) in 50 mM ammonium bicarbonate (A6141 Sigma-Aldrich), pH 8.0, incubated at 75°C for 1 h. For both culture media and whole spheroid samples, disulfide bonds were reduced by the addition of dithiothreitol (D9779, Sigma-Aldrich) to a final concentration of 5 mM, followed by incubation at 56°C for 30 min. Free sulfhydryl groups were alkylated with 10 mM iodoacetamide (I1149, Sigma-Aldrich) for 30 min at room temperature in the dark. Proteins were digested with trypsin (V5280, Promega) at 37°C overnight in a thermocycler, and the reaction was quenched by the addition of 0.5% (v/v) formic acid (28905, Thermofisher Scientific).

Resulting peptide samples were loaded onto Evotip Pure disposable trap columns (Evosep Biosystem) according to the manufacturer’s protocol. Peptide separation was performed on an Evosep One liquid chromatography system (Evosep Biosystems) coupled online to a timsTOF HT mass spectrometer (Bruker Daltonics) equipped with a CaptiveSpray nano-electrospray ionisation source. Peptide separation was achieved on an analytical column packed with C18 reversed-phase material. The timsTOF HT was operated in data-independent acquisition (DIA) mode using the dia-PASEF acquisition scheme, as previously described[8]. Raw dia-PASEF data files were processed using Spectronaut software (version 20.0; Biognosys AG) with the directDIA analysis pipeline. Spectral libraries were generated *in silico* from human protein sequences from the Uniprot database (https://www.uniprot.org/).

Peptide and protein identifications were filtered at a false discovery rate (FDR) of 1% at both the precursor and protein group levels using the target-decoy strategy in Spectronaut. Ion mobility information (1/K0 values) was used for scoring to improve identification confidence. Quantification was performed at the MS2 level using the integrated peak areas of fragment ions, and cross-run normalisation was applied using the global median strategy. Protein group quantities were calculated using the MaxLFQ algorithm as implemented in Spectronaut[9]. Data matrices were exported from Spectronaut for downstream statistical analysis.

**Additional information relating to differentially expressed protein analysis**

Proteomic data were analysed by the University of Edinburgh's Discovery Research Platform for Hidden Cell Biology Bioinformatics Core. Normalised MaxLFQ protein abundances were used for differential protein enrichment analysis, performed in R using the limma[10] and DEP[11] packages. Proteins were filtered to retain those that had more than one hit present in all replicates of at least one condition. Missing values were imputed using the QRILC method. Proteins were considered significantly differentially expressed if they met both a threshold of an adjusted p-value $<0.05$ following Benjamini-Hochberg false discovery rate correction and a minimum $\log_2$FC of $\pm0.25$.

#### Supplemental Figure 1 Wittke, Watt et al.

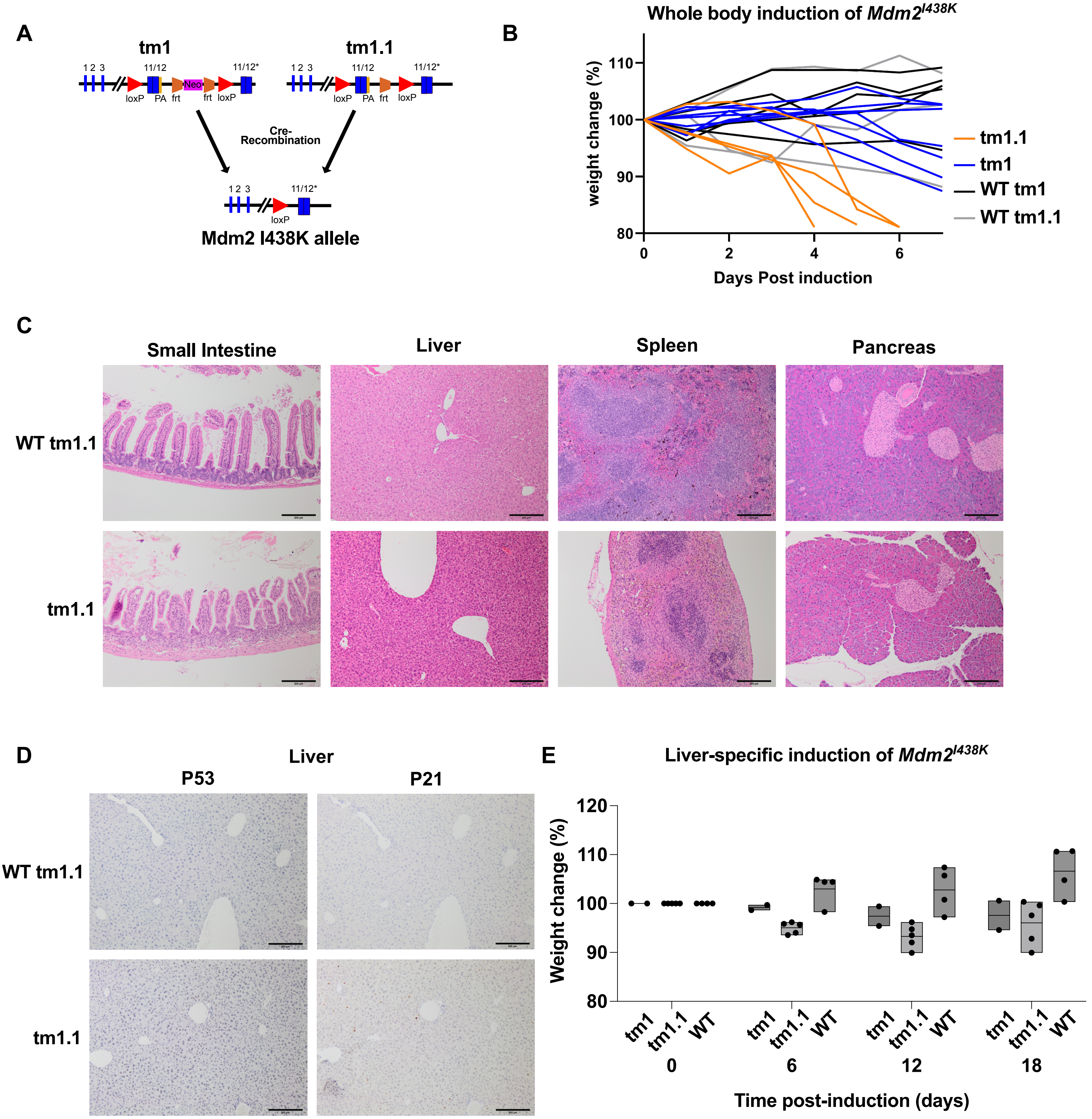


**Supplemental Figure 1: Whole-body induction of *Mdm2*$^{I438K}$ in adult mice has divergent strain and allele-specific consequences**

A: Comparison between the two versions of the *Mdm2*$^{I438K}$ allele (tm1 and tm1.1) used in this study. The *Mdm2*$^{I438K}$ tm1 allele contains *Mdm2* WT exons 1-12, with exons 11/12 flanked by loxP sites to allow for Cre-mediated excision. Downstream, a construct containing WT exon 11 and I438K mutant exon 12 (12*) is not expressed in unrecombined cells. The tm1 allele retains an FRT-flanked Neomycin (Neo) selection cassette between WT exons 11/12 and the second loxP site that was utilised for ES cell selection. To create the tm1.1 strain, the Neo selection cassette was removed by FLP-FRT-mediated recombination, resulting in *Mdm2*$^{I438K}$ mice harbouring the tm1.1 allele. Cre-mediated recombination of either the tm1 or tm1.1 allele allows for expression of *Mdm2*$^{I438K}$ in recombined tissues. This recombination product is identical between tm1 and tm1.1 mice.

B: Weight change (%) following gene induction (days) in tm1 and tm1.1 *Mdm2*$^{I438K}$ mice or *Mdm2*$^{WT}$ controls following *Rosa*-Cre$^{ER}$-mediated whole-body induction of the allele. N=7 tm1 mice (N=4 male, N=3 female), N=4 tm1.1 mice (N=1 male, N=3 female), N=5 WT mice of the same strain background as tm1 mice (WT-tm1) (N=3 male, N=2 female), and N=3 mice of the same strain background as tm1.1 mice (WT-tm1.1) (N=1 male, N=2 female). All tm1.1 mice reached the defined 20% weight-loss threshold for clinical endpoint between 4-6 days after induction.

C: Representative H&E images of small intestine, liver, spleen, and pancreas from tm1.1 mice at clinical endpoint or WT mice from (B). N-numbers as in (B). Scale bars 200 μm.

D: Representative p53 and p21 staining of the liver of tm1.1 mice at clinical endpoint or WT mice from (B). N=2 tm1.1 mice and N=3 WT mice. The other N=2 tm1.1 mice from (B) were not sampled for liver tissue. Scale bars 200 μm.

E: Weight change (%) following AAV8-TBG-Cre mediated liver-specific induction of the *Mdm2*$^{I438K}$ allele (days) in tm1 and tm1.1 *Mdm2*$^{I438K}$ mice or *Mdm2*$^{WT}$ controls (WT). N=2 tm1 mice (N=2 male, N=0 female), N=5 tm1.1 mice (N=3 male, N=2 female), and N=4 WT mice comprising a mix of tm1 background mice (N=2 male, N=0 female) and tm1.1 background mice (N=1 male, N=1 female). Data points are from individual mice and presented as mean +/- range for each timepoint.

#### Supplemental Figure 2 Wittke, Watt et al.

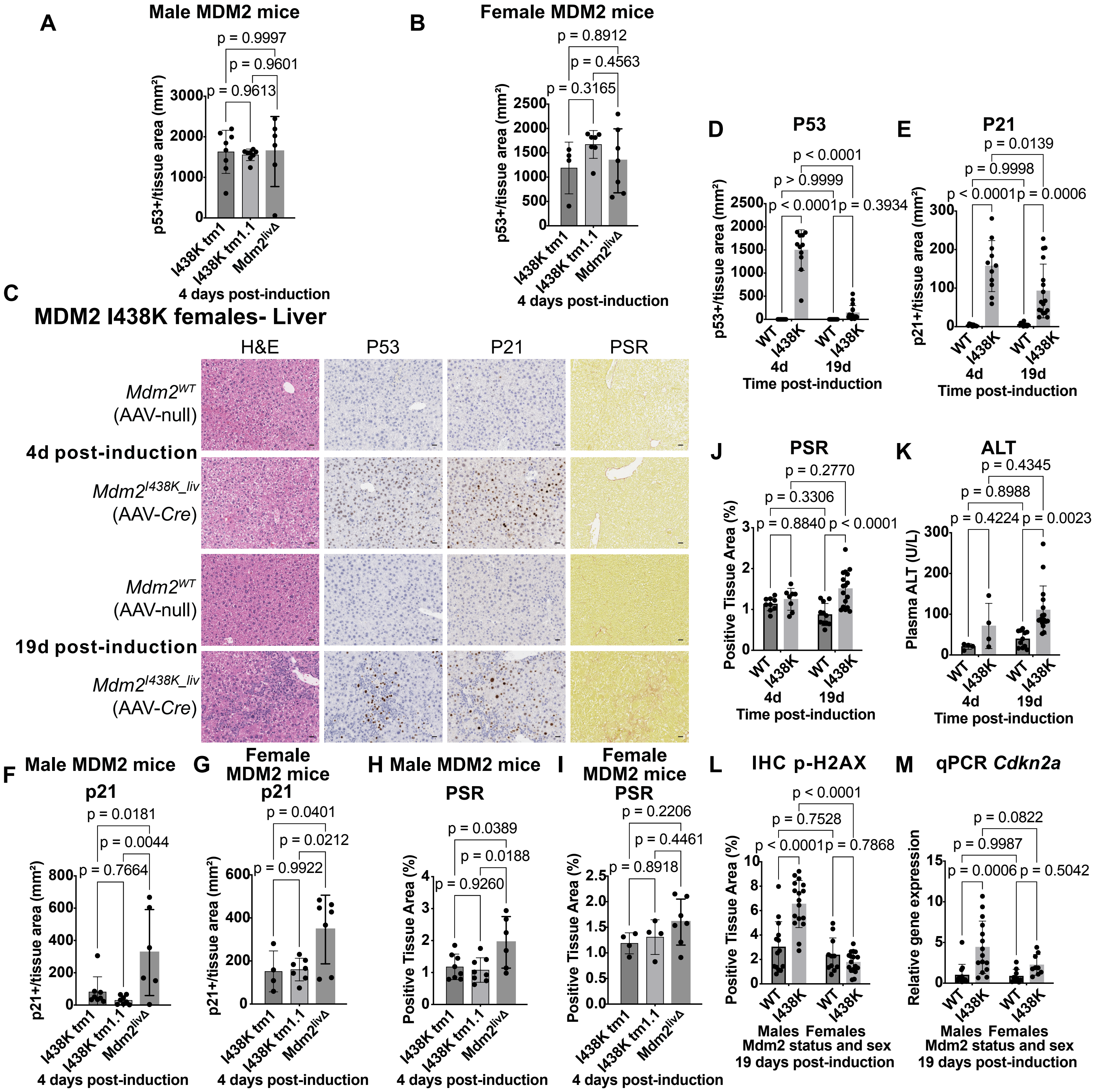


**Supplemental Figure 2: The hepatocellular response to loss of MDM2 E3 activity is similar across $Mdm2^{I438K}$ alleles but differs between male and female $Mdm2^{I438K\_liv}$ mice**

A/B: Comparison of p53-positive hepatocytes per tissue area ($mm^2$) in male (A) and female (B) AAV8-TBG-Cre induced tm1 and tm1.1 strains of $Mdm2^{I438K}$ mice (I438K tm1 and I438K tm1.1) and $Mdm2^{Ex5/6\_liv\Delta}$ mice ($Mdm2^{liv\Delta}$) at 4 days post-induction. N=8 I438K tm1 and I438K tm1.1 male mice and N=6 male $Mdm2^{liv\Delta}$ male mice. N=4 I438K tm1 female mice, N=7 I438K tm1.1 female mice, and N=7 $Mdm2^{liv\Delta}$ female mice. These tm1 and tm1.1 samples are pooled for results presented in Figure 1 (males) and Figure S2 (females). Data points are from individual mice and analysed using a one-way ANOVA with Tukey's multiple comparisons test. Multiplicity-adjusted p-values as shown. Bars show mean +/- SD.

C: Representative liver H&E images and staining for p53, p21, and Picro Sirius Red (PSR) in female AAV8-TBG-Cre induced $Mdm2^{I438K}$ mice ($Mdm2^{I438K\_liv}$) and control AAV8-TBG-Null-treated $Mdm2^{I438K}$ mice ($Mdm2^{WT}$) at 4 or 19 days post-induction. Data are pooled between tm1 and tm1.1 $Mdm2^{I438K}$ strains in each condition. At 4d, images are representative of N=9 $Mdm2^{WT}$ mice and N=11 $Mdm2^{I438K\_liv}$ mice. At 19d, images are representative of N=11 $Mdm2^{WT}$ mice and N=17 $Mdm2^{I438K\_liv}$ mice. Scale bars 20 μm.

D/E: Quantification of p53 (D) or p21 (E)-positive hepatocytes per tissue area ($mm^2$) in female mice from (C). WT: $Mdm2^{WT}$. N-numbers as in (C). Data points are from individual mice and analysed using a one-way ANOVA with Tukey's multiple comparisons test. Multiplicity-adjusted p-values as shown. Bars show mean +/- SD.

F/G: Comparison of p21-positive hepatocytes per tissue area ($mm^2$) in male (F) and female (G) mice at 4 days post-induction as in A/B. N-numbers, data presentation, and analysis as in A/B. These tm1 and tm1.1 samples are pooled for results presented in Figure 1 (males) and Figure S2 (females).

H/I: Comparison of PSR positive tissue area (%) in male (H) and female (I) mice at 4 days post-induction as in A/B. N=8 I438K tm1 and I438K tm1.1 male mice and N=6 male $Mdm2^{liv\Delta}$ male mice. N=4 I438K tm1 female mice, N=4 I438K tm1.1 female mice, and N=7 $Mdm2^{liv\Delta}$ female mice. Data presented and analysed as in A/B.

J: Quantification of PSR positive tissue area (%) in female mice from (C). At 4d, N=9 $Mdm2^{WT}$ mice and N=8 $Mdm2^{I438K\_liv}$ mice. Samples from N=3 $Mdm2^{I438K\_liv}$ mice at 4d were inadvertently not stained for PSR. At 19d, N=11 $Mdm2^{WT}$ mice and N=17 $Mdm2^{I438K\_liv}$ mice.

K: Quantification of Alanine Transaminase (ALT) activity (U/L) in plasma samples from female mice in (C). At 4d, N=4 $Mdm2^{WT}$ and $Mdm2^{I438K\_liv}$ mice. At 19d, N=11 $Mdm2^{WT}$ mice and N=16 $Mdm2^{I438K\_liv}$ mice. Samples were not obtained or analysed for N=5 $Mdm2^{WT}$ or N=5 $Mdm2^{I438K\_liv}$ 4d mice or from N=1 $Mdm2^{I438K\_liv}$ 19d mouse.

L: IHC quantification of phospho-H2AX (Ser139) (p-H2AX) positive tissue area in male and female $Mdm2^{I438K\_liv}$ and $Mdm2^{WT}$ mice at 19 days post-induction. N=14 male $Mdm2^{WT}$ mice, N=17 male $Mdm2^{I438K\_liv}$ mice, N=11 female $Mdm2^{WT}$ mice, and N=17 female $Mdm2^{I438K\_liv}$ mice.

M: RT-qPCR analysis of liver *Cdkn2a* expression relative to *Actin* in male and female $Mdm2^{I438K\_liv}$ and $Mdm2^{WT}$ mice at 19d post-induction. N=13 $Mdm2^{WT}$ and N=15 $Mdm2^{I438K\_liv}$ male mice and N=9 female mice per group.

For graphs in (J-M), data points are from individual mice and analysed using a two-way ANOVA with Tukey's multiple comparisons test. Multiplicity-adjusted p-values as shown. Bars show mean +/- SD.

**Supplemental Figure 3 Wittke, Watt et al.**

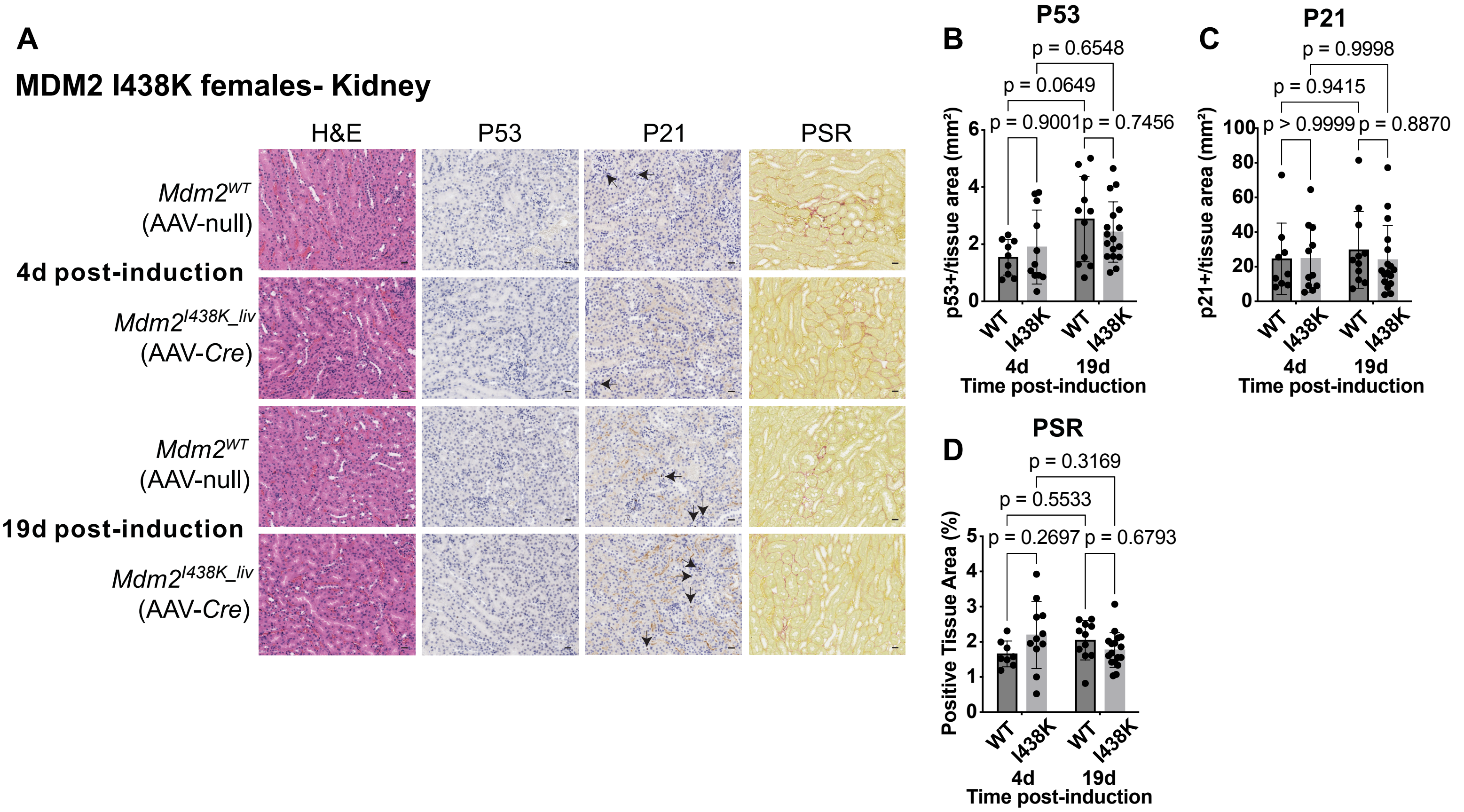


**Supplemental Figure 3: Loss of liver MDM2 E3 activity does not alter p21 or fibrosis in the kidneys of female Mdm2$^{I438K_liv}$ mice**
A: Representative kidney H&E images and staining for p53, p21, and PSR in female *Mdm2*$^{I438K\_liv}$ and control *Mdm2*$^{WT}$ mice at 4 or 19 days post-induction. Data are pooled between tm1 and tm1.1 *Mdm2*$^{I438K}$ strains in each condition. Images are focused on the cortex and include the proximal tubule compartment. At 4d, N=9 *Mdm2*$^{WT}$ mice and N=11 *Mdm2*$^{I438K\_liv}$ mice. At 19d, N=11 *Mdm2*$^{WT}$ mice and N=17 *Mdm2*$^{I438K\_liv}$ mice. Arrows denote p21-positive cells. Scale bars 20 µm.
B-D: Quantification of p53 (B) or p21 (C)-positive cells per tissue area (mm$^2$), and PSR positive tissue area (%) (D) in mice from (A). WT: *Mdm2*$^{WT}$. N-numbers as in (A). Data points are from individual mice and analysed using a two-way ANOVA with Tukey's multiple comparisons test. Multiplicity-adjusted p-values as shown. Bars show mean +/- SD.

### Supplemental Figure 4 Wittke, Watt et al.

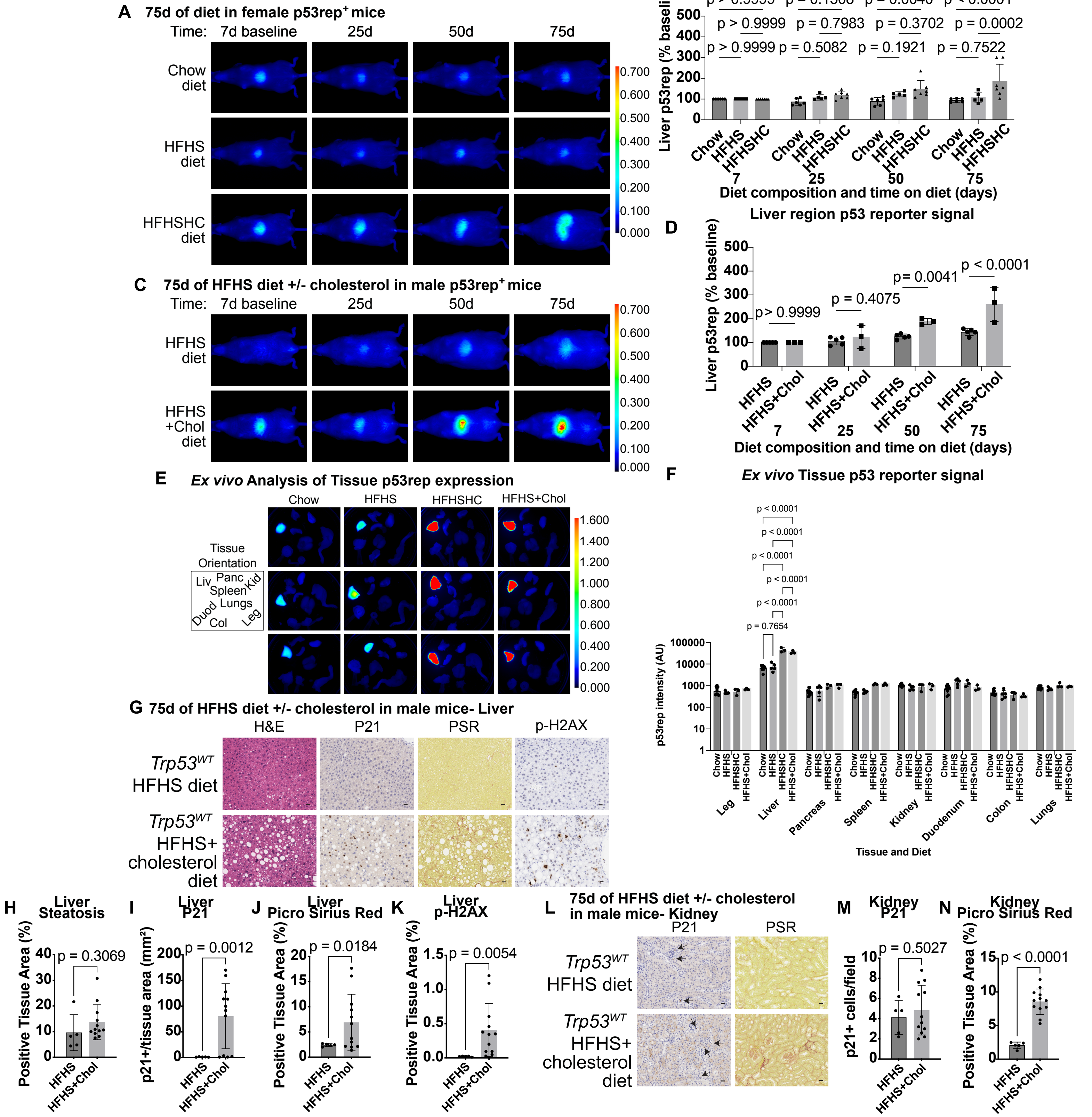


**Supplemental Figure 4: Cholesterol promotes robust liver p53 activation and MASH *in vivo***

A/B: Female whole-body p53 reporter (p53rep) mice were given obesogenic high fat high sugar (HFHS) diet or high fat high sugar high cholesterol (HFHSHC) diet at 65-75 days of age or remained on normal chow diet and imaged after the indicated days. Representative images (A) and p53rep signal quantification (B) normalised to the liver signal identified per mouse at the baseline (7d diet) measurement. N=6 chow, N=5 HFHS, and N=7 HFHSHC mice imaged per timepoint. LUT intensity values for images in (A) as shown.

C/D: Male p53rep mice were given obesogenic high fat high sugar (HFHS) diet or HFHS diet supplemented with high cholesterol (HFHS+Chol) diet at 65-75 days of age. Representative images (C) and P53rep signal quantification (D) normalised to the liver signal identified per mouse at the baseline (7d diet) measurement. N=5 HFHS, and N=3 HFHS+Chol mice imaged per timepoint. Data from HFHS-fed mice are the same as presented in Figure 3A/B. Images are also duplicated for consistency of comparison. LUT intensity values for images in (C) as shown.

E/F: Representative images (E) and quantification (arbitrary units (AU)) (F) of *ex vivo* p53Rep signal in tissues from male reporter mice after 75 days on HFHS, HFHSHC, HFHS+Chol, or chow control diet. N=10 chow, N=5 HFHS, N=3 HFHSHC, and N=3 HFHS+Chol samples. Tissue orientation in scanned images and LUT intensity profile as shown for (E). Data presented on a log scale for (F).

For graphs in (A-F), data points are from individual mice and analysed using a two-way ANOVA with Tukey's multiple comparisons test. Multiplicity-adjusted p-values as shown. Bars show mean +/- SD.

G: Representative liver H&E images and staining for p21, p-H2AX, and PSR in *Trp53* WT male mice given HFHS or HFHS+Chol diet for 75 days. N=5 HFHS mice and N=12 HFHS+Chol mice. Representative images used for HFHS mice are duplicated from Figure 3C for consistency of comparison. Scale bars 20 µm.

H-K: Quantification of steatosis area (%) in H&E slides (H), p21-positive cells per tissue area (mm²) (I), or PSR (J) or p-H2AX (K)-positive tissue area (%) in mice from (G). N-numbers as in (G). Data from HFHS-fed mice are the same as presented in Figure 3 D-G.

L: Representative kidney staining for p21 and PSR in male mice from (G). HFHS-fed cohort mice are duplicated from Figure 3. Images used for HFHS mice are duplicated from Figure 3H for consistency of comparison. Images are focused on the cortex and include the proximal tubule compartment. N=5 HFHS mice and N=12 HFHS+Chol mice. Arrows denote p21-positive cells. Scale bars 20 µm.

M/N: Quantification of p21-positive cells per field of view (M) and PSR positive tissue area (%) (N) in mice from (L). Data from HFHS-fed mice are the same as presented in Figure 3 H-J.

For graphs in (H-N), data points are from individual mice and analysed using an unpaired two-tailed T-test with Welch's correction. P-value as shown. Bars show mean +/- SD.

#### Supplemental Figure 5 Wittke, Watt et al.

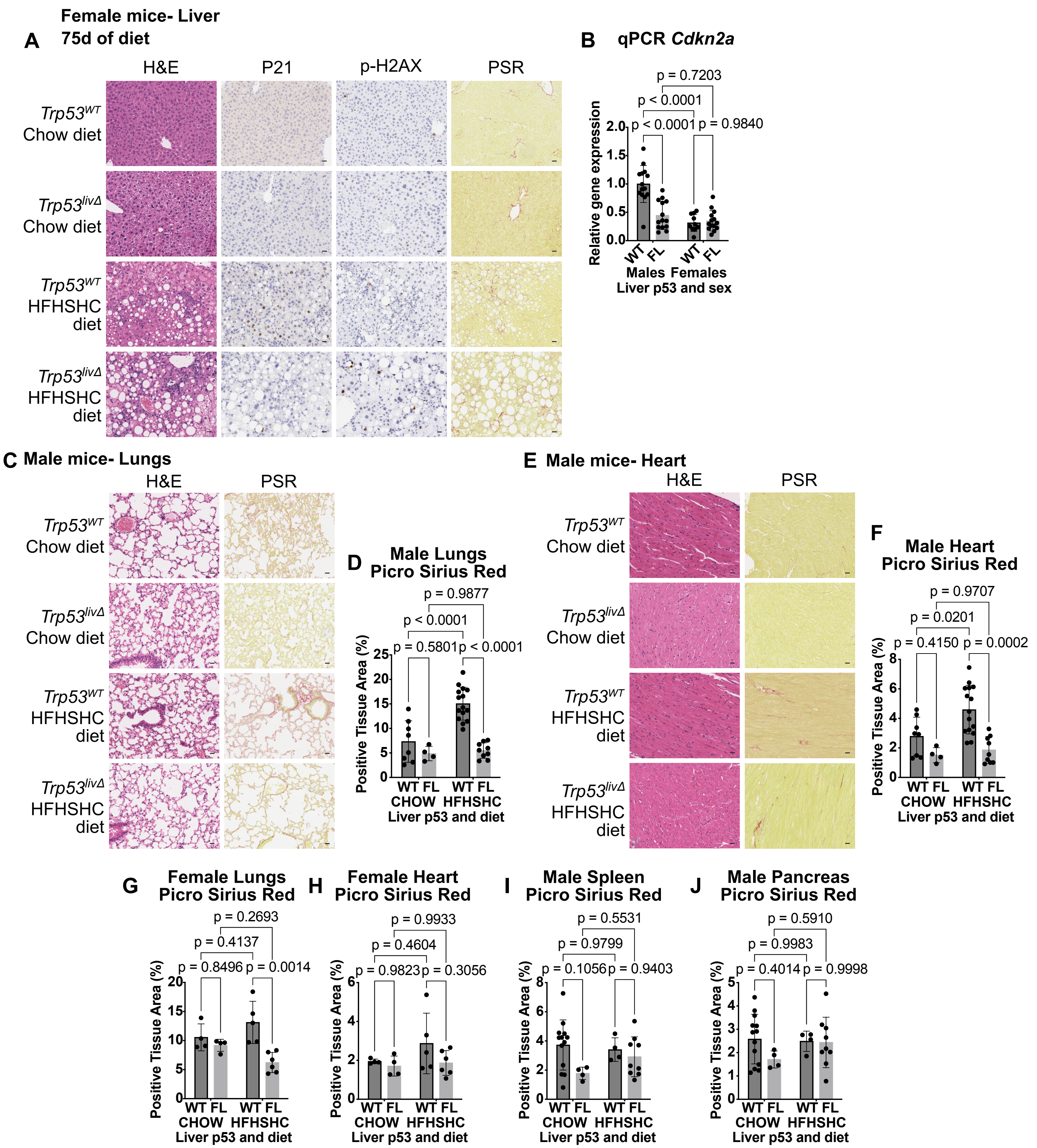


**Supplemental Figure 5: Hepatic p53 enhances liver fibrosis in MASH and is necessary for distal induction of p21 and fibrosis *in vivo***

A: Representative liver H&E images and staining for p21, p-H2AX, and PSR in female AAV8-TBG-Cre induced *Trp53*$^{FL/FL}$ (*Trp53*$^{liv\Delta}$) mice or *Trp53* wildtype (*Trp53*$^{WT}$) mice given HFHSHC or control chow diet for 75 days. N=9 *Trp53*$^{WT}$ chow mice, N=9 *Trp53*$^{liv\Delta}$ chow mice, N=10 *Trp53*$^{WT}$ HFHSHC mice and N=15 *Trp53*$^{liv\Delta}$ HFHSHC mice. *Trp53*$^{WT}$ mice comprised a mix of AAV8-TBG-Cre-treated *Trp53* WT mice and uninduced *Trp53*$^{FL/FL}$ mice and include both p53rep$^{+}$ and p53rep$^{-}$ mice. For further information, see supplementary methods. Scale bars 20 µm.

B: RT-qPCR analysis of liver *Cdkn2a* expression relative to *Actin* in male and female *Trp53*$^{liv\Delta}$ and *Trp53*$^{WT}$ mice given HFHSHC diet for 75 days. N=14 male mice per condition, N=10 *Trp53*$^{WT}$ female mice and N=14 *Trp53*$^{liv\Delta}$ female mice.

C: Representative lung H&E images and staining for PSR in male *Trp53*$^{liv\Delta}$ or *Trp53*$^{WT}$ mice given HFHSHC or control chow diet for 75 days. N=8 *Trp53*$^{WT}$ chow mice, N=4 *Trp53*$^{liv\Delta}$ chow mice, N=15 *Trp53*$^{WT}$ HFHSHC mice, and N=9 *Trp53*$^{liv\Delta}$ HFHSHC mice. Scale bars 20 µm.

D: Quantification of PSR positive tissue area (%) in mice from (C). N-numbers as in (C).

E/F: Representative images of heart H&E and staining for PSR (E) and quantification of PSR positive tissue area (%) (F) in male *Trp53*$^{WT}$ mice given HFHSHC or control chow diet for 75 days. Scale bars 20 µm. N-numbers as in (C).

G/H: Quantification of PSR positive tissue area (%) in the lungs (G) and heart (H) of female *Trp53*$^{liv\Delta}$ or *Trp53*$^{WT}$ mice given HFHSHC or control chow diet for 75 days as in (A). N=4 chow mice per genotype, N=5 *Trp53*$^{WT}$ HFHSHC-fed mice and N=6 *Trp53*$^{liv\Delta}$ HFHSHC-fed mice.

I/J: Quantification of PSR positive tissue area (%) in the spleen (I) and pancreas (J) of male *Trp53*$^{liv\Delta}$ and *Trp53*$^{WT}$ mice given HFHSHC diet for 75 days. N=13 *Trp53*$^{WT}$ chow mice, N=4 *Trp53*$^{liv\Delta}$ chow mice, N=4 *Trp53*$^{WT}$ HFHSHC mice, and N=9 *Trp53*$^{liv\Delta}$ HFHSHC mice.

For all graphs, data points are from individual mice and analysed using a two-way ANOVA with Tukey's multiple comparisons test. Multiplicity-adjusted p-values as shown. Bars show mean +/- SD.

### Supplemental Figure 6 Wittke, Watt et al.

**A** Circulating proteins differentially present in serum after 75d HFHSHC diet

| Chow - HFHSHC | Mean difference | Discovery? | q value | Individual P Value |
|---|---|---|---|---|
| GDF-15 | -1.7090 | Yes | 0.0360 | 0.0028 |
| LDLR | -1.7090 | Yes | 0.0360 | 0.0028 |
| Serpin E1 | -1.6860 | Yes | 0.0360 | 0.0032 |
| DPPIV/CD26 | -1.6390 | Yes | 0.0360 | 0.0042 |
| IL-12 | -1.6340 | Yes | 0.0360 | 0.0043 |
| Osteoprotegerin (TNFRSF11B) | -1.6210 | Yes | 0.0360 | 0.0046 |
| BAFF/BLyS/TNFSF13B | -1.6190 | Yes | 0.0360 | 0.0047 |
| Pentraxin 3 | -1.5890 | Yes | 0.0360 | 0.0055 |
| CXCL10/IP-10 | -1.5880 | Yes | 0.0360 | 0.0055 |
| CD14 | -1.5770 | Yes | 0.0360 | 0.0058 |
| IL-7 | -1.5750 | Yes | 0.0360 | 0.0059 |
| Leptin | -1.5740 | Yes | 0.0360 | 0.0060 |
| Angiopoietin-2 | -1.5440 | Yes | 0.0360 | 0.0069 |
| IL-27 | -1.5220 | Yes | 0.0360 | 0.0078 |
| PCSK9 | -1.5190 | Yes | 0.0360 | 0.0079 |
| IL-11 | -1.4830 | Yes | 0.0360 | 0.0095 |
| IL-33 | -1.4820 | Yes | 0.0360 | 0.0096 |
| E-Selectin | -1.4460 | Yes | 0.0414 | 0.0115 |
| Gas 6 | -1.4050 | Yes | 0.0487 | 0.0140 |
| MMP-3 | 1.4900 | Yes | 0.0360 | 0.0092 |
| Flt-3 Ligand | 1.4950 | Yes | 0.0360 | 0.0089 |
| FGF acidic (FGF1) | 1.5150 | Yes | 0.0360 | 0.0081 |
| CX3CL1 | 1.5200 | Yes | 0.0360 | 0.0079 |
| Reg3G | 1.5280 | Yes | 0.0360 | 0.0076 |
| RAGE | 1.6550 | Yes | 0.0360 | 0.0038 |
| Chemerin | 1.6880 | Yes | 0.0360 | 0.0032 |

**B** Murine Circulating Hits analysed within MASH Human Plasma Proteomics Govaere *et al.* 2023

| F0-F2 - F3/F4 | Mean difference | Discovery? | q value | Individual P Value |
|---|---|---|---|---|
| SELE | -1.1130 | Yes | <0.0001 | <0.0001 |
| TNFRSF11B | -0.7896 | Yes | <0.0001 | <0.0001 |
| GDF15 | -0.7557 | Yes | <0.0001 | <0.0001 |
| ANGPT2 | -0.4632 | Yes | 0.0046 | 0.0014 |
| CD14 | -0.4618 | Yes | 0.0046 | 0.0014 |
| TNFSF13B | -0.4601 | Yes | 0.0046 | 0.0015 |
| CXCL10 | -0.3108 | No | 0.0664 | 0.0316 |
| GAS6 | -0.2856 | No | 0.0912 | 0.0483 |
| FLT3LG | -0.2052 | No | 0.2676 | 0.1558 |
| IL12 | -0.1840 | No | 0.2952 | 0.2030 |
| LDLR | -0.1618 | No | 0.3554 | 0.2632 |
| SERPINE1 | -0.1554 | No | 0.3559 | 0.2825 |
| LEP | -0.1082 | No | 0.4472 | 0.4542 |
| CX3CL1 | -0.1037 | No | 0.4472 | 0.4732 |
| IL7 | -0.0244 | No | 0.7115 | 0.8659 |
| MMP3 | -0.0142 | No | 0.7261 | 0.9220 |
| REG3G | 0.0048 | No | 0.7360 | 0.9735 |
| DPP4 | 0.0429 | No | 0.6585 | 0.7665 |
| IL11 | 0.0759 | No | 0.5397 | 0.5997 |
| RARRES2 | 0.1268 | No | 0.3995 | 0.3805 |
| FGF1 | 0.1291 | No | 0.3995 | 0.3719 |
| PTX3 | 0.1389 | No | 0.3977 | 0.3367 |
| IL27 | 0.1882 | No | 0.2952 | 0.1930 |
| AGER | 0.3212 | No | 0.0622 | 0.0263 |
| PCSK9 | 0.4358 | Yes | 0.0070 | 0.0026 |

**C**

**Proteomic mass spectrometry:**
**Control vs early MASH (LPON) LiverACE human spheroids**

| LPON vs. Control | | Whole spheroid proteomics | | | Culture media secretomics | | |
|---|---|---|---|---|---|---|---|
| Protein | Protein ID | p val | adj. p val | log2 FC | p val | adj. p val | log2 FC |
| SERPINE1 | P05121 | 0.00000008 | 0.00000825 | 1.11000000 | 0.00835159 | 0.05850000 | 0.29400000 |
| LDLR | P01130 | 0.00019811 | 0.00296000 | 0.23700000 | nd | | |
| GDF15 | Q99988 | 0.01298297 | 0.05960000 | 0.46500000 | 0.52425345 | 0.73300000 | 0.23500000 |
| RARRES2 | Q99969 | 0.02098250 | 0.08300000 | 0.24400000 | 0.01475822 | 0.08790000 | 0.25300000 |
| IL27RA | Q6UWB1 | 0.07756293 | 0.20100000 | 0.15200000 | nd | | |
| CD14 | P08571 | 0.16161548 | 0.32500000 | -0.10900000 | nd | | |
| DPP4 | P27487 | 0.20978563 | 0.38600000 | 0.14100000 | 0.79606056 | 0.90100000 | 0.02210000 |
| AGER | Q15109 | nd | | | nd | | |
| ANGPT2 | O15123 | nd | | | nd | | |
| CX3CL1 | P78423 | nd | | | nd | | |
| CXCL10 | P02778 | nd | | | nd | | |
| FGF1 | P05230 | nd | | | nd | | |
| FLT3LG | P49771 | nd | | | nd | | |
| GAS6 | Q14393 | nd | | | nd | | |
| IL11 | P20809 | nd | | | nd | | |
| IL12A | P29459 | nd | | | nd | | |
| IL12B | P29460 | nd | | | nd | | |
| IL27A | Q8NEV9 | nd | | | nd | | |
| IL27B | Q14213 | nd | | | nd | | |
| IL7 | P13232 | nd | | | nd | | |
| LEP | P41159 | nd | | | nd | | |
| MMP3 | Q13368 | nd | | | nd | | |
| PCSK9 | Q8NBP7 | nd | | | nd | | |
| PTX3 | P26022 | nd | | | nd | | |
| REG3G | Q6UW15 | nd | | | nd | | |
| SELE | P16581 | nd | | | nd | | |
| TNFRSF11B | O00300 | nd | | | nd | | |
| TNFSF13B | Q9Y275 | nd | | | nd | | |

**Human RNA-seq- MASH**

**GSE130970**

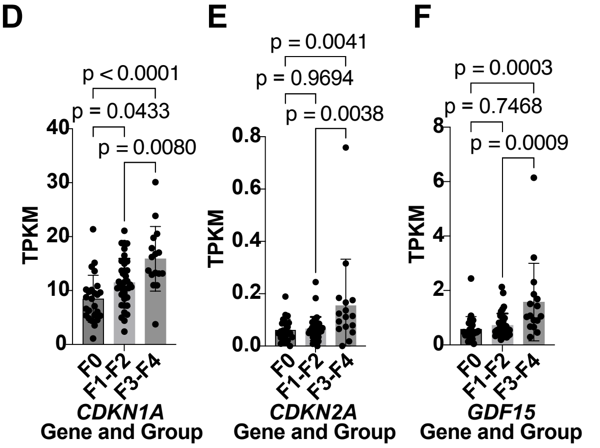


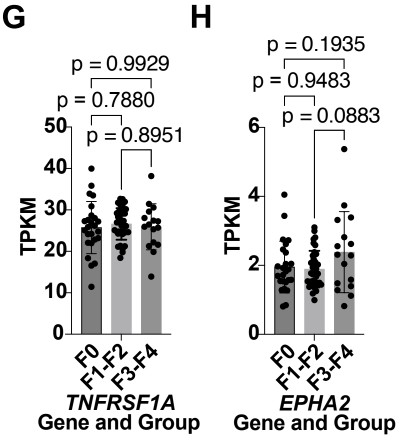


**Human RNA-seq- MASH**

**GSE135251**

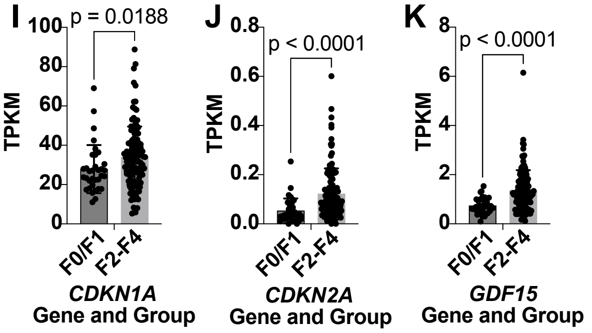


**Supplemental Figure 6: GDF15 is induced by hepatic p53 and correlates with markers of fibrotic kidney damage in humans**

A: List of the circulating proteins differentially present in the serum of male mice given HFHSHC or control chow diet for 75 days. Z-score normalised values for all 111 targets per Proteome Profiler Mouse XL Cytokine Array were analysed using a two-way ANOVA with the two-stage linear step-up procedure of Benjamini, Krieger and Yekutieli to correct for multiple comparisons by controlling FDR at q < 0.05. Circulating proteins in the list met this threshold and are ranked by mean difference. Mean difference values given as (Chow – HFHSHC). N=4 mice per condition. Q-value and individual p-values per circulating protein as indicated.

B: List of the significant circulating proteins identified in (A) within the published plasma proteomics dataset from human MASH patients reported by Govaere and colleagues in 2023. In the Govaere *et al.* dataset, 25/26 significant circulating proteins from (A) were detected and compared between patients with advanced MASH (F3/F4 fibrosis score) and those with less advanced disease (F0-F2 fibrosis score). Further information on fibrosis score in Materials and Methods. N=112 F0-F2 and N=79 F3/F4 samples. Q-value and individual p-values per circulating protein as indicated.

C: Whole spheroid proteomic mass spectrometry (proteomics) and mass spectrometry of cell culture media (secretomics) from human hepatic acinar (LiverACE) spheroids. LiverACE spheroids were cultured in medium supplemented with lactate, pyruvate, octanoic acid, and ammonium chloride (LPON) for 72 h to induce steatosis and features of early MASH or remained in baseline medium (control). The relative abundance of the 26 circulating proteins identified in (A) were examined within both datasets. Two proteins (IL12 and IL27) were assessed for both A and B variants, leading to N=28 samples assessed. Of these 28, 7 were detected in the proteomics dataset and 4 were detected in the secretomics dataset. Proteins presented in rank order based on proteomic p-value. Differential proteins were required to meet both the Benjamini–Hochberg-adjusted p<0.05 and |log2FC|≥0.25 criteria. 'nd'- not detected. For spheroid proteomics, N=10 control and N=11 LPON samples. For spheroid secretomics, N=15 samples per condition. P-value, adjusted p-value, and fold change per protein as shown. Further information available in Supplementary Methods.

D-H: Analysis of *CDKN1A* (D), *CDKN2A* (E), *GDF15* (F), *TNFRSF1A* (G) and *EPHA2* (H) liver gene expression in the public transcriptomics dataset GSE130970 comparing MASH patients of low fibrosis score (F0), moderate fibrosis score (F1/F2), and high fibrosis score (F3-F4). N=25 F0, N=37 F1/F2, and N=16 F3/F4 samples. Data points are from individual patients and analysed using a one-way ANOVA with Tukey's multiple comparisons test. Multiplicity-adjusted p-values as shown. Bars show mean +/- SD.

I-K: Analysis of *CDKN1A* (I), *CDKN2A* (J), and *GDF15* (K) liver gene expression in the public transcriptomics dataset GSE135251 comparing MASH patients of low fibrosis score (F0/F1) and MASH patients with high fibrosis score (F2-F4). N=34 F0/F1 and N=121 F2-F4 samples. Data presented in transcripts per kilobase million (TPKM). Data points are from individual patients and analysed using an unpaired two-tailed T-test with Welch's correction. P-value as shown. Bars show mean +/- SD.
